# Non-specific vs specific DNA binding free energetics of a transcription factor domain protein for target search and recognition

**DOI:** 10.1101/2022.12.14.520393

**Authors:** Carmen Al Masri, Biao Wan, Jin Yu

## Abstract

Transcription factor (TF) proteins regulate gene expression by binding to specific sites on the genome. In the facilitated diffusion model, an optimized search process is achieved by the TF protein alternating between 3D diffusion in the bulk and 1D diffusion along DNA. While undergoing 1D diffusion, the protein can switch from a search mode for fast diffusion along non-specific DNA to a recognition mode for stable binding to specific DNA. It was recently noticed that for a small TF domain protein, re-orientations on DNA other than conformational changes happen between the non-specific and specific DNA binding. We here conducted all-atom molecular dynamics simulations with steering forces to reveal the protein-DNA binding free energetics, with a difference between the non-specific and specific binding about 10 *k*_*B*_*T*, confirming that the search and recognition modes are distinguished only by protein orientations on the DNA. As the binding free energy difference differs from that being estimated from experimental measurements about 4-5 *k*_*B*_*T* on 15-bp DNA constructs, we hypothesize that the discrepancy comes from DNA sequences flanking the 6-bp central binding sites impacting on the dissociation kinetics measurements. The hypothesis is supported by a simplified spherical protein-DNA model along with stochastic simulations and kinetic modeling.

## Introduction

A key component in regulation of gene expression is the network of proteins known as transcription factors (TFs), which recognize and bind to specific DNA sequences known as transcription factor binding sites (TFBS), usually 6–20 bp long with variable sequences [1]. Once bound to the TFBS, TFs can either promote or block the recruitment of other proteins and enzymes that initiate transcription (e.g, RNA polymerase), which results in genes being either activated or silenced [1, 2]. It is therefore essential to understand how TFs can efficiently navigate through the genome, finding their target binding sites among millions to billions of base pairs [3], at a rate that is up to about 100 times faster than what would be predicted by free diffusion or random collisions of the TF with a target site on the genome [4].

A widely accepted theoretical framework for this search process is the facilitated diffusion model [4, 5, 6]. In this model, the protein diffuses in three-dimensional (3D) space until it randomly collides with a DNA site, which is most likely a non-specific binding site. The TF afterwards can slide along the DNA as keeping contact with it, or hop from one site on the DNA to the next. Such a phenomenon is regarded as the one-dimensional (1D) diffusion. The TF thus alternates between the 1D and 3D diffusion processes until it reaches the target site on the genome. This model has been supported by various studies from both *in vitro* [7, 8, 9, 10, 11, 12] and *in vivo* [13, 14, 15, 16, 17].

The 1D diffusion has been further modeled by considering the interaction potential energy between the protein and DNA to be sequence-dependent [18]. The studies estimate the roughness of the diffusion free energy landscape, which was shown to have a fluctuation magnitude of 1-2 *k*_*B*_*T*. On the other hand, once the TF protein locates the target site, it is able to form a stable protein-DNA complex. Such a stability appears to require the binding free energy amplitude to be much larger than *k*_*B*_*T* [18], which is seemingly incompatible with the fast search requirement. To resolve the paradox, it was proposed that the protein adopts two conformations (referred as modes or states) while bound to the DNA: one is the search mode, with relatively weak protein-DNA interactions to allow for fast protein 1D diffusion, and the other is the recognition mode, which happens when the protein locates the target site, supporting the formation of a stabilized protein-DNA complex. On the other hand, a recent all-atom molecular dynamics (MD) simulation study on a small TF domain protein WRKY has shown that the protein conformation change between nonspecific (corresponding to search mode) and specific (recognition mode) DNA binding is not necessary [19]. Starting from a crystal structure of the WRKY domain TF bound to a specific DNA sequence (i.e, the W-box), the DNA sequence was subsequently changed to nonspecific or poly-A sequences. Without notable protein conformational changes, an onset of diffusion and a 1-base pair (bp) diffusional stepping of WRKY were captured on the non-specific and the poly-A DNA, respectively, within the 10*μ*s of equilibrium MD simulations. In contrast, WRKY remained stably bound to the specific DNA sequence within the control simulations. Given that the protein conformations in the three systems were almost identical while the protein orientations on the DNA varied significantly, it was therefore suggested that the protein orientational changes with respect to the DNA rather than the conformational changes are sufficient to lower the protein-DNA interaction energies, i.e., for the protein to switch between non-specific (search) and specific (recognition) modes. Nevertheless, calculations on protein-DNA binding free energetics to substantially distinguish the search and recognition modes have not been conducted yet. For the non-specific DNA binding, a possibility remains that the necessary protein conformational change for switching from the recognition mode, to the search mode has not yet been sampled within the MD simulations, such that the protein might remain in the recognition mode upon initial binding to the non-specific DNA.

Accordingly, in current work, we aim at substantially distinguishing the protein recognition on the specific DNA and the search on the non-specific DNA by calculating the binding free energetics of the WRKY domain protein on the specific and nonspecific DNA, respectively (Figure 1). It is expected that the protein stably bound on the specific DNA in the recognition mode would display a highly stabilized binding free energy 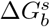 (with 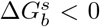 and 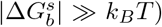), while the protein in association with the non-specific DNA in the search mode would show a significantly smaller magnitude of the binding free energy 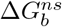 (with 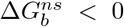 and 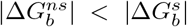). For this purpose, we started from the microsecond equilibrated WRKY domain protein structures bound with specific and non-specific DNA from the previous studies [19, 20], and calculated the protein binding free energy energetics of 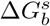 and 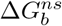, respectively, by enforcing dissociation of the protein from the DNA binding site, in the all-atom MD simulations. The binding free energy Δ*G*_*b*_ was obtained using two methods: the Jarzynski’s equality [21] and then the umbrella sampling [22], both widely used for free energy calculations [23, 24, 25]. We confirmed that although the protein conformations in the two systems remained identical, as the protein orientations on the DNA vary significantly, the resulting binding free energies become well distinguished, i.e., 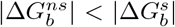 well holds. Meanwhile, we also examined the protein dissociation dynamics during the enforced protein-DNA dissociation processes. It appears that the dissociation well couples with the protein diffusion or horizontal motions along the DNA and with substantial protein orientational changes, as hydrogen bonding interactions initially formed at the protein-DNA interface gradually disappear during the enforced dissociation. Finally, we compared the calculated protein binding free energy difference between the specific and nonspecific DNA ΔΔ*G*_*b*_ with that being evaluated according to experimentally measured dissociation constants. A discrepancy was revealed, which we hypothesized to be due to impacts from DNA sequences flanking to the central DNA binding site. Such a proposal was then investigated through a simplified spherical protein model along with stochastic dynamics simulations, which consistently show that the flanking DNA regions affect protein dissociation kinetics so that to interfere with measuring protein-DNA binding affinities upon varying DNA sequences at the central DNA binding site.

**Figure 1:**
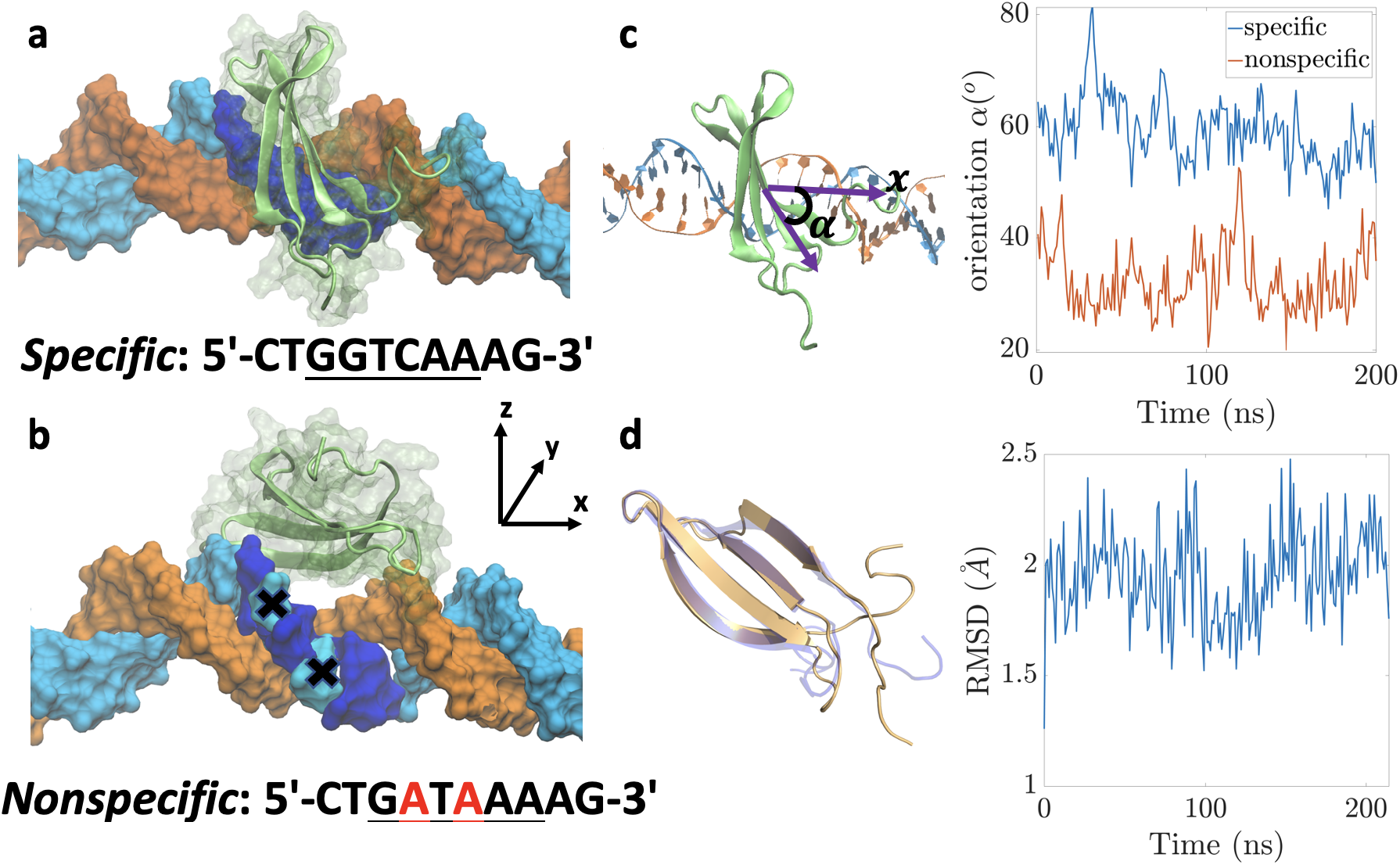
An overview of the MD simulation systems. **(a), (b)** show the WRKY domain protein bound to the specific (GGTCAA) and nonspecific (GATAAA) sequence, respectively. The binding site is shown in dark blue, with the mutated residues in the nonspecific system in cyan labeled with an X. The preferred strand is shown in cyan, non-preferred in orange. **(c)** shows the protein orientations on the DNA with respect to the x-axis (DNA long axis) for specific and nonspecific systems, with the orientation angle *α* defined as shown in the left diagram. **(d)** shows the difference in protein conformations between specific and non-specific systems is shown, with the left figure showing the two protein conformations aligned. The protein from the specific system is shown in transparent blue, and that from the nonspecific one in orange. The proteins were aligned using pyMOL by excluding outlier atoms over 5 iterations and the RMSD between the two structures was computed. The graph shows the RMSD difference between the two proteins throughout the equilibration (frame by frame comparison), averaging around 1.95 ± 0.2 Å.

## Materials and Methods

### System setup

To compare specific and non-specific DNA binding of the WRKY domain protein, we simulated correspondingly the two systems: the WRKY1-N in complex with the W-box DNA obtained from crystal structure (accession PDB code 6J4E) and the WRKY1-N in complex with a mutated W-box sequence. We refer to the former as the specific DNA (5’-CT<u>GGTCAA</u>AG-3’) and to the latter as the nonspecific DNA (5’-CT<u>GATAAA</u>AG-3’). We followed the protocol in [19]: to avoid DNA end-effects in the simulations, the 15-bp DNA construct obtained from the crystal structure was extended to 34 bp by adding the same random nucleotides to the two sides of the specific and nonspecific DNA. The CYS and HIS residues in the WRKY protein around a zinc ion were changed to CYM (deprotonated state) and HIE (residue 164, with *γ*-nitrogen protonated) or HID (residues 138 and 166, with *ϵ*-nitrogen protonated), respectively, to form the stable zinc finger domain. For starting structures for our current all-atom MD simulations, we took snapshots at *t* = 4*μ*s from a previously generated 10-*μ*s equilibrium trajectories for both specific and nonspecific DNA binding systems [19]. The simulation systems are shown in Figure 1a,b.

### MD simulations setup

All-atom MD simulations were performed with GROMACS 2020.4 [26] under the Amber99SB-ILDN force field [27] for protein and the Parmbsc1 (BCS1) force field for nucleic acids [28]. For both systems, the protein-DNA complex was solvated with TIP3P water in a rectangular box, with a minimum distance from the complex to the boundary of the simulation box of 15 Å. The system was neutralized with Na^+^ and Cl^−^ ions at an ionic concentration of ~ 0.15 M. The total simulation system consists of ~ 85,000 atoms. Periodic boundary conditions were applied. The van der Waals (vdW) and short-range electrostatic interactions used a cutoff of 10 Å. The particle-mesh Ewald (PME) method was applied to deal with the long-range electrostatic interactions [29].

The solvated system was minimized with the steepest-descent algorithm, followed by a 4-ns equilibration under the NVT ensemble then a 4-ns equilibration under the NPT ensemble with a time step of 2 fs. Position restraints with a force constant of 1000 kJ.mol^−1^nm^−2^ were imposed on the heavy atoms of the system. The simulations were run using the leap-frog stochastic dynamics integrator. The temperature was set to 300 K and the inverse friction constant was set to 2 ps. The pressure was set at 1 bar using the Parrinello-Rahman Barostat pressure control method [30]. Finally, the position restraints were lifted and a final equilibration of 200 ns was conducted, with restraints applied to the edges of the DNA only (heavy atoms of the first and last residues; with a force constant of 1000 kJ.mol^−1^nm^−2^) to keep the DNA axis aligned to the x-axis.

### Equilibrium reached for the specific and nonspecific complexes

To make sure equilibrium has been reached in the MD simulations for both specific and non-specific systems, the RMSD of the protein as well as its orientation and radius of gyration changes were examined. From the final 200 ns unconstrained equilibration, we find that after ~ 80 ns, the RMSD of the protein conformation is stabilized for both the specific and nonspecific systems (Figure S1a). We also find that the protein in the nonspecific system has a more significant orientation change over time (16.6° ±7.9° for nonspecific vs 10.1° ± 2.8° for specific) (Figure S1b). Throughout the equilibration process, the protein orientation angles with respect to the DNA *α* were measured to be quite different between the specific (57 ° ± 8°) and nonspecific (32° ± 3°) systems (Figure 1c). We also checked the change in the radius of gyration over time and found it to be small (−0.06 ± 0.16 Å for the specific system vs −0.25 ± 0.19 Å for the nonspecific system) (Figure S1c), again indicating that the protein conformation has reached equilibrium in both systems.

### Steered MD (SMD) simulations setup

To force the dissociation of the protein, its center of mass (COM) was steered along a reaction coordinate *ξ*, which was taken along the vertical dissociation path with respect to the DNA (oriented along the x-axis). In the current coordinate system, it was found that the − *y* direction for the specific DNA system and +*z* direction for the nonspecific DNA system have the protein dissociation most ready or probable (Figure 2, bottom), as demonstrated by randomly steering the the protein COM using the RAMD implementation in GROMACS [31] (Figure 2, top). A spring force with a force constant of 3000 kJ.mol^−1^nm^−2^ was applied to the COM of the protein, steering it along *ξ* at a speed of 0.1 Å/ns. To avoid the DNA following along the protein while pulling, the heavy atoms of the DNA were restrained by harmonic restoring forces with a force constant of 1000 kJ.mol^−1^nm^−2^.

**Figure 2:**
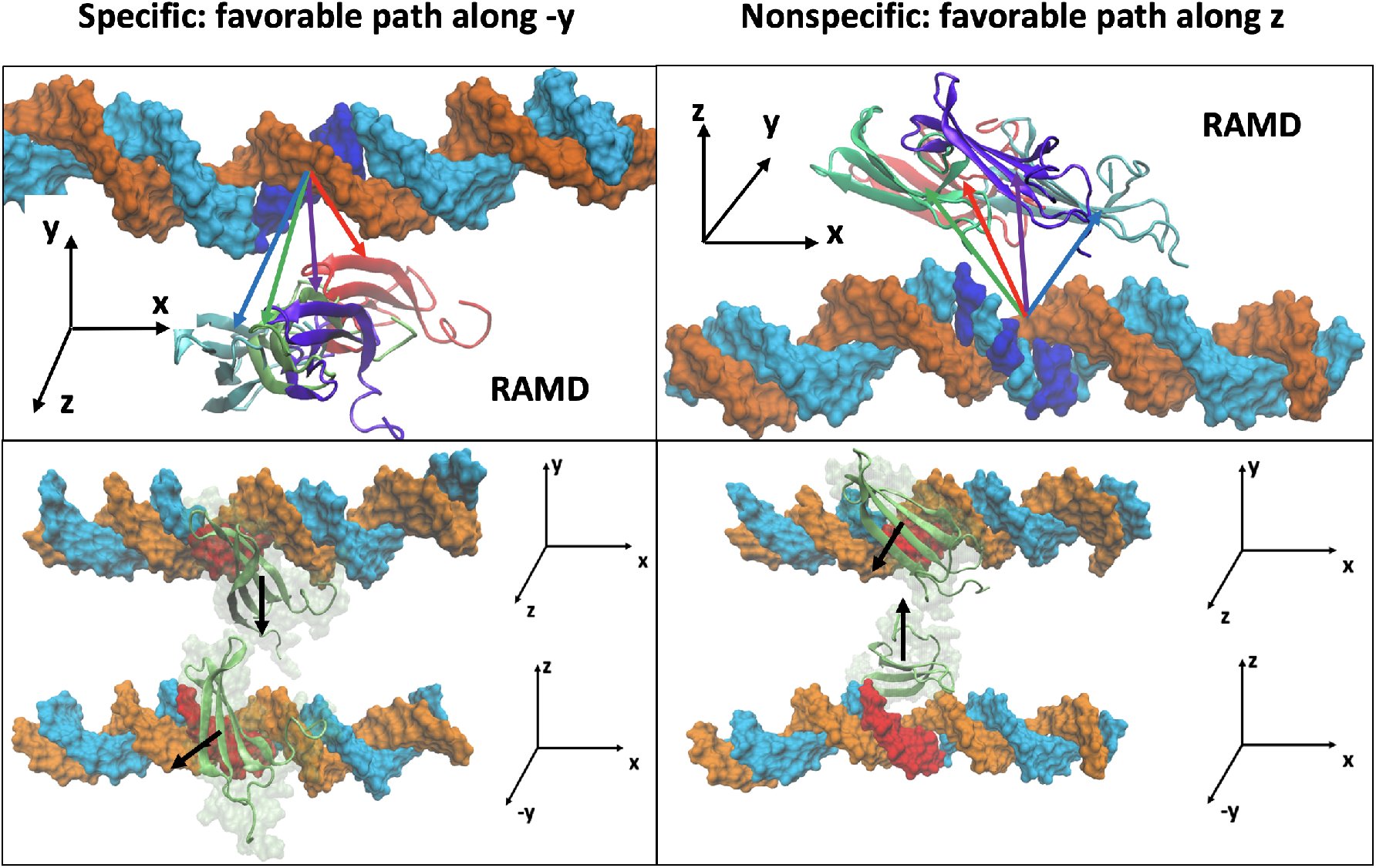
Top: Choosing directions of the protein dissociation for both specific and nonspecific systems, by employing Randomly Accelerated MD (RAMD): the COM of the protein is steered in a random direction with a constant force k=2000 kJ.mol^−1^nm^−2^. If the protein COM does not move by a distance larger than 0.25 Å in 100 fs, the direction is updated to another random one until the protein reaches a maximum distance set by the user (2nm for current purposes). A snapshot at the time of dissociation is shown for 4 different trajectories in both specific (left) and nonspecific (right) DNA binding systems, with the arrows pointing towards four dominant directions of the protein random dissociation (protein and arrows colored by corresponding trajectory sampled). Bottom: The preferred direction of the protein dissociation shown from different perspectives for each system. It was found that the − *y*-direction is preferred in the specific binding system and the *z*-direction in the nonspecific one. Details and supporting figures can be found in the SI and Figure S2.

### Potential of Mean Force (PMF) calculations using the Jarzynski’s equality

Jarzynski’s equality [21] enables one to obtain free energy differences from the work done through non-quasi static processes. The equality is given by:

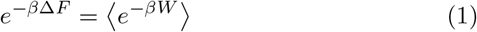

where Δ*F* is the free energy difference, *W* is the work, ⟨…⟩ denotes the ensemble average and *β* = 1*/k*_*B*_*T*, with *T* the temperature of the system. The main concern of the equality is that the exponential is dominated by small work values that rarely get sampled, making the equality computationally expensive to implement. This difficulty can be sometimes reduced by using its approximate form via cumulant expansion [21], assuming that the work distribution satisfies certain forms, such as the Gaussian distribution. Using the Jarzynski’s equality, one can then compute the free energy profile as a function of the reaction coordinate Φ(*ξ*) (or the PMF). The path along *ξ* was sampled via the SMD simulations [32], using a guiding harmonic potential to constrain *ξ* to follow a parameter *λ* as detailed in the above section. In the stiff-spring approximation, the expression for the PMF was found to be [33]:

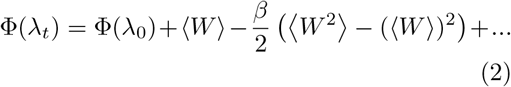

where *λ*_*t*_ and *λ*_0_ are the values of the parameter *λ* at times t and 0, and *W* is the work done on the system sampled in the time interval between 0 and *t*. If the distribution of work is Gaussian (i.e., with the stiff spring approximation), the third and higher cumulants all vanish.

To reduce biases in the sampling, we implemented the following procedures: five starting frames were taken from the last of the 200-ns NPT ensemble equilibrium simulations to generate SMD trajectories. Similar to what has been implemented in previous studies [34, 35], the pulling was conducted in short segments of length 1 Å for *ξ* ≤ 4 Å, then of length 4 Å for *ξ* ≥ 4 Å, alternating between forward and reverse pulling events and allowing the two neighboring segments share the initial positions or overlap for the final positions (within a region of 0.5 Å). Before pulling, the initial structure for each segment of the pulling simulation was equilibrated for 2 ns. The scheme and more details can be found in the SI and Figure S3.

### Calculations using umbrella sampling

The free energy along a reaction coordinate *ξ* (or the PMF) is directly related to how *ξ* is populated. Umbrella sampling is a popular method that allows the sampling of energetically unfavorable regions along *ξ* by applying an external potential and restraining the system near the given region, allowing for the PMF Φ(*ξ*) to be computed [22]. More specifically, sampling of *ξ* is enhanced near a certain region *λ* by applying a harmonic position restraint. A set of *N* simulations were launched along the reaction coordinate *ξ*, each biasing the system near the region parametrized by *λ*_*i*_ with *i* = 1, …, *N*. We refer to the sets of simulations as umbrella sampling windows. In each of these windows, the distribution of *ξ* will be recorded as the umbrella histogram *h*_*i*_(*ξ*), which represents the biased probability distribution 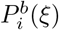 along *ξ*. The PMF is then obtained by reweighting or unbiasing the probability distributions from the umbrella sampling. A widely used way to conduct the unbiasing is the weighted histogram analysis method (WHAM) [36]. For our purposes, we use the GROMACS implementation of WHAM g wham [37] to get the unbiased probability distribution *P* (*ξ*), from which we can obtain the PMF through the relation:

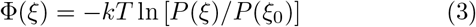

where *P* (*ξ*_0_) is the unbiased probability distribution of the arbitrary reference point *ξ*_0_. To sample a path as described in the “SMD simulations setup” section, a single SMD trajectory pulling the protein away from DNA with a duration of 200 ns was launched. This procedure makes it possible for the protein to dissociate at around 100 ns, as evident from the complete breaking of hydrogen bonds (HBs) (Figure S4).

For both the specific and nonspecific DNA-binding systems, 21 umbrella sampling windows along the trajectory were then launched, such that the spacing between the centers of two neighboring window is about 0.1 nm. The first umbrella window was taken at *ξ* = −0.1 nm from the starting position to improve the sampling around that initial protein-DNA bound configuration.

Each window was then run for 200 ns with the COM of the protein restrained by a harmonic potential using a force constant of 3000 kJ.mol^−1^nm^−2^ and the sampling was used to generate a PMF for the protein-DNA dissociation. Finally, the convergence of the PMFs was examined (Figure S5). To estimate the errors along the obtained PMF, Bayesian bootstrapping of histograms was conducted as detailed in [37] whereby each umbrella window generated 10 independent histograms, which were then chosen by sampling with replacement to generate a new PMF. Each histogram has then a corresponding probability to be sampled given by a random weight. Finally, the average PMF is obtained, with the errors being the standard deviations along the bootstrapped PMFs.

### Simplified spherical protein model

In order to investigate how the protein binding free energetics obtained from the protein-DNA dissociation PMFs connect with the protein dissociation events and kinetics sampled randomly, in a highly simplified quantitative model, the domain protein is modeled by a charged sphere (Figure S6). In this model, discretized positive and negative charges were placed on the surface of the spherical protein with overall neutrality. The DNA is modeled by a line of charges. The protein-binding sites on the DNA were then represented by linearly aligned points with spacing of 0.34 nm. In addition, the protein-DNA interfacial HB interactions were represented by the pair-interaction between the sites on the protein hemisphere and the sites on DNA. The protein-DNA electrostatic interactions screened by solution with ions can be approximated by the Debye-Huckel potential, and the interfacial HB interaction can be described by the Morse potential depending on the bond angle or protein orientation. More details on the implementation can be found in the SI (section S-V, Figure S6).

## Results

To verify whether the protein-DNA binding free energies for the non-specific and specific DNA constructs are consistent with the definitions of search and recognition modes, respectively, we first computed the PMFs using the Jarzynski’s equality for both systems, derived the respective binding free energies, and then estimated the difference between the binding free energies of the two systems. Given that the SMD trajectories showed limited samplings of some of the protein’s degrees of freedom, we proceeded to compute the PMFs using the umbrella sampling method. After that, we investigated causes of discrepancies between computationally determined binding free energy differences and experimentally measured protein-DNA binding affinities by suggesting potential effects of flanking sequences on the binding affinity measurements using a highly simplified spherical protein model. In such a model, we also computed PMFs of protein dissociation from DNA and measured protein dissociation kinetics, using steered and spontaneous Langevin dynamics simulations, respectively.

### Binding free energy measurements using SMD & Jarzynski’s equality

In order to obtain the binding free energetics of the WRKY domain protein in both specific and non-specific DNA binding systems, we implemented first the Jarzynski’s equality to calculate the PMF of pulling the protein away from the DNA. 10 trajectories for each system were generated and utilized for data analyses. The reaction coordinate *ξ* was defined as the distance over which the protein was pulled, with the pulling path detailed in Figure 2. The work applied as a function of *ξ* for all 20 trajectories sampled is displayed in Figure 3a. As expected, we find that the average total work done to the specific system is larger than that of the nonspecific one (50 ± 14 *k*_*B*_*T* vs 35 ± 8 *k*_*B*_*T*). Meanwhile, the work distribution had a high variance for the specific and the nonspecific systems, and deviated significantly from a Gaussian (Figure S7a,b). Due to the high fluctuations, convergence of the PMFs obtained by taking only the first two cumulants of the work distributions is not well achieved (Eq (2), Figure S8a,b). Consequently, the PMFs are thus not well behaved (Figure S8c).

**Figure 3:**
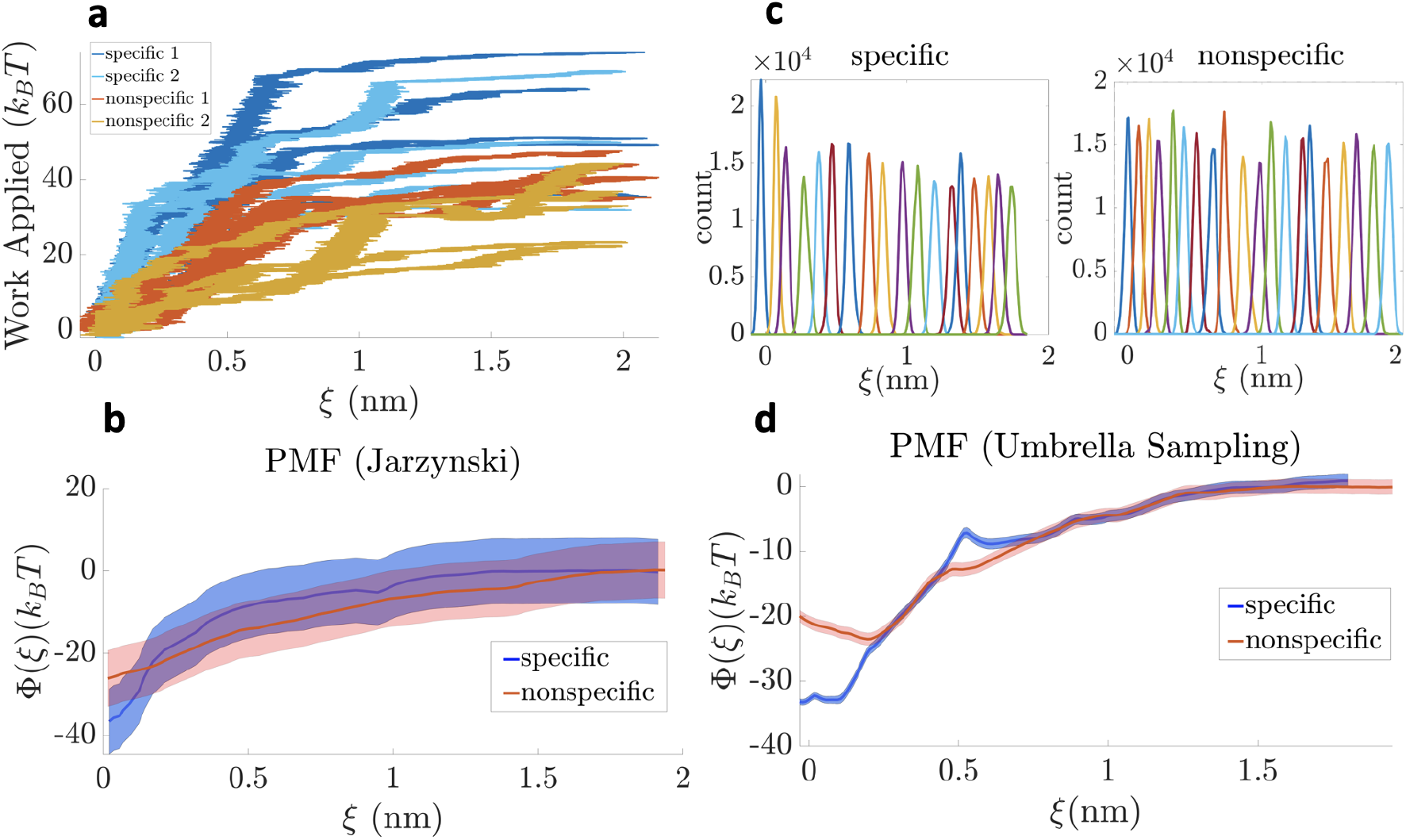
PMFs of enforced protein dissociation from DNA from implementing both Jarzynski’s equality and umbrella sampling methods. **(a)** shows the work from all SMD trajectories for both specific (blue) and nonspecific (orange) systems. Dark colors correspond to setup 1 and light colors to setup 2, as detailed in Supplementary section S-III. **(b)** shows the resulting PMFs obtained from Jarzynski’s equality. **(c)** shows the distributions of *ξ*, the reaction coordinate of protein pulling, in each umbrella window for both specific and nonspecific DNA-binding systems, and **(d)** the resulting PMFs from umbrella sampling. The standard deviations are shown as the shaded regions.

For improved results, we constructed the PMF utilizing the original exponential averaging form of the Eq (1). The convergence on the obtained PMF then shows (Figure S8d,e), i.e, within 4 *k*_*B*_*T* for specific and 2 *k*_*B*_*T* for nonspecific binding systems. On the PMFs shown (Figure 3b), one finds that the protein dissociation along *ξ* demonstrates a higher energy barrier in the specific system than that in the nonspecific.

One can then obtain the binding free energy Δ*G*_*b*_ from the PMF Φ(*ξ*) as Δ*G*_*b*_ = Φ(*ξ*_0_) − Φ(*ξ*_∞_), where Φ(*ξ*_0_) is the free energy of the inital protein-DNA bound state, and Φ(*ξ*_∞_) is the free energy after protein dissociation from the DNA, which we can set to zero as a reference value. Accordingly, we obtain 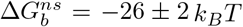 and 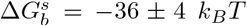, corresponding to the nonspecific and specific binding free energies, respectively. The binding free energy difference between the systems is then given by 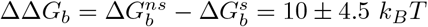.

Finally, we checked whether the trajectories sufficiently sample the protein translational and rotational degrees of freedom (Figure 4a). We find that horizontal diffusional displacements of the protein are well sampled. Interestingly, the protein in the specific system showed more significant horizontal diffusional motions than in the non-specific system, likely due to a larger dissociation barrier in the specific system that restrains the protein from vertical displacements away from the DNA. Nevertheless, the protein does not reorient much throughout the trajectory, implying insufficient sampling of the protein rotational degrees of freedom.

**Figure 4:**
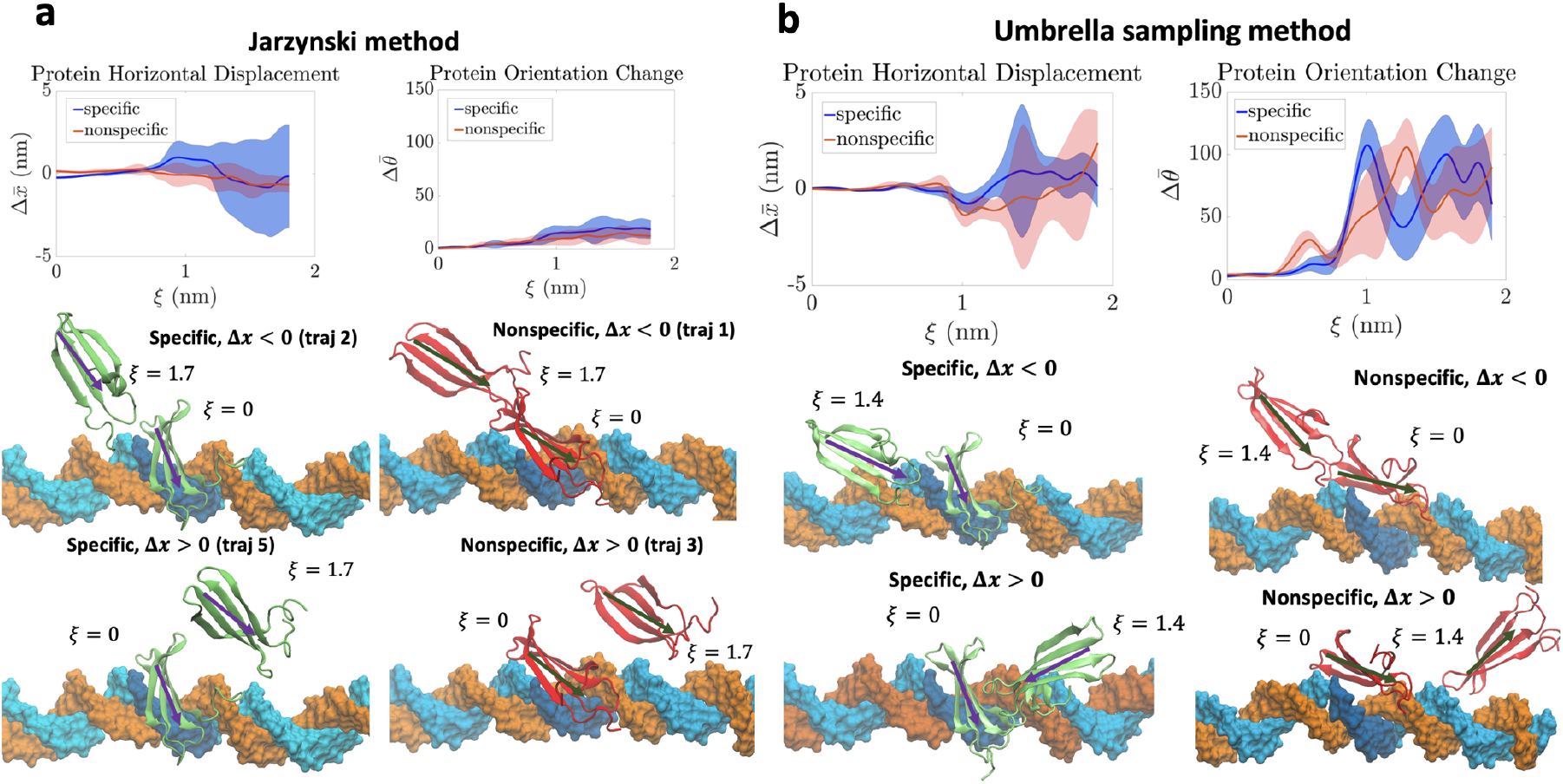
Protein horizontal displacements and orientational changes during enforced dissociation for construction of PMFs using Jarzynski’s method **(a)** and umbrella sampling **(b)**. Each of the top left panels in **(a)** and **(b)** show the protein horizontal displacements along the DNA long axis, Δ*x*, and the top right panels the protein orientation changes, Δ*θ*. The standard deviation of each quantity is shown as the shaded region. Each of the bottom panels show snapshots taken at the initial bound state (*ξ* = 0) and the dissociated state (*ξ* = 1.7 in the Jarzynski method and *ξ* = 1.4 in the umbrella sampling method). The protein bound to the specific sequence is colored in green and that bound to the nonspecific sequence in red. After dissociating, the protein can diffuse either to the left (Δ*x <* 0, upper panels) or to the right (Δ*x >* 0, lower panels) in both specific and nonspecific constructs. Note that the range of orientation changes is more limited in the Jarzynski method than the umbrella sampling.

### Binding free energy measurements using Umbrella Sampling

PMFs of the WRKY protein dissociation from DNA were then also constructed using the umbrella sampling method, by launching 20 umbrella windows along the vertical dissociation path *ξ* for the protein to be away from DNA. At each window, 200 ns MD simulation with the protein center constrained at a particular distance *ξ* was conducted, and the convergence of the resulted PMF was examined over time (Figure S5). The convergence was then observed around 150 ns for both specific and nonspecific protein-DNA binding systems, with the PMF fluctuating within 1.5 *k*_*B*_*T*, of the order of thermal fluctuations. Accordingly, the last 50 ns of simulation data for each window were considered for further analyses.

The corresponding histograms at 200 ns are shown in Figure 3c, and the final PMFs obtained along with the error from the Bayesian bootstrapping analyses are shown in Figure 3d. From the PMF results, we get the protein-DNA binding free energies 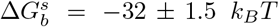 and 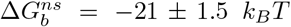 for the specific and non-specific systems, respectively. The difference in the binding free energies between the two systems is then ΔΔ*G*_*b*_ = 11 ± 2 *k*_*B*_*T*. Comparing with the PMF calculations from the Jarzynski’s method, the energetic fluctuations become smaller in the umbrella sampling implementation. The average value of the binding free energy difference between the non-specific and specific DNA binding system of the WRKY shows consistently at ~ 10 *k*_*B*_*T*.

Since it was experimentally measured that the dissociation constants *K*_*D*_ = *k*_*off*_ */k*_*on*_ for the specific and nonspecific binding systems are respectively as 0.1 *μM* and 8 *μM* [19], the corre-sponding binding free energy difference would be estimated as ΔΔ*G*_*b*_ = ln 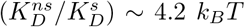. However, both the PMF calculations above indicate that the binding free energy difference are ΔΔ*G*_*b*_ ~ 10 *k*_*B*_*T*. in those experiments, however, 15-bp DNA constructs were used for such measurements [19, 20]. To resolve the discrepancy, we probed in the later section whether the flanking sequences included in DNA construct impact on the protein dissociation measurements.

### Protein dissociation dynamics features near DNA

To probe additional structural dynamics or relaxation of the protein during the enforced dissociation, we calculated the changes in the horizontal displacements or diffusion of the protein along the DNA and the orientational changes of the protein with respect to the DNA (Figure 4a,b, top) along the DNA dissociation direction *ξ*, for both specific and nonspecific binding systems. We find that in both methods, after an initial stage of dissociation (*ξ >* 1 nm), as the protein-DNA interfacial HBs are largely broken, the protein starts diffusing comparatively freely along the DNA long axis (the x-direction). In addition, with sufficient sampling, the protein rotational degrees of freedom become notable upon dissociation. One can see that the orientation changes of the protein are indeed much better sampled in the umbrella sampling method (up to 120 degrees of protein spatial rotations; see Figure 4b, upper right panel) than in the Jarzynski equality method (Figure 4a, upper right panel). The improved sampling on the protein rotational degrees of freedom may contribute to reduced energetic fluctuations in the PMFs calculated above using the umbrella sampling method. It is worth noting that in the nonspecific system the protein starts rotating as early as *ξ* = 0.5 nm, due to the protein being more weakly bound to the non-specific DNA and having a lower dissociation energy barrier than to the specific DNA. In comparison, in the specific binding system, the protein starts freely rotating at *ξ* ≈1 nm. Meanwhile, the horizontal diffusion of the protein becomes significant in the dissociation (*ξ* ≈ 1.5 nm; Figure 4b, upper left) for the specific DNA binding earlier than for the nonspecific DNA binding, which is also due to the higher barrier to hinder the protein dissociation from the specific DNA. To gain more insights into the protein dissociation dynamics, the HB occupancy and dynamics at the protein-DNA association interface were evaluated from the umbrella sampling simulations (Figure 5), which allow sufficient sampling of the protein rotational relaxation. We find that the specific protein-DNA binding system forms HB contacts mostly on the specific sequence on one DNA strand (or the preferred strand; 5’-CT<u>GGTCAA</u>AG-3’), which are abruptly broken upon protein dissociation at ~ 1 nm. After the abrupt dissociation, very few protein-DNA HB contacts re-form (up to ~ 8 new HB contacts). Additionally, only a few residues form HB contacts with the other (i.e., non-preferred) strand throughout the dissociation process. In contrast, in the nonspecific system, the protein dissociates gradually starting at ~ 0.5 nm, with initially formed HBs broken, while new and weak HB contacts emerge thereafter (up to ∵35 new HB contacts). Additionally, at *ξ* = 0−0.1 nm, we find that the HB contacts with the preferred strand are significantly reduced in the non-specific DNA binding system, in comparison with the specific DNA binding system. Upon the protein dissociation initiated at *ξ* ~ 0.5 nm, there appears to be no strand preference with non-specific DNA, as residues form HB contacts with both DNA strands almost equally (3 ± 1 HBs in the originally preferred strand and 2 ± 2 HBs in the originally non-preferred strand).

**Figure 5:**
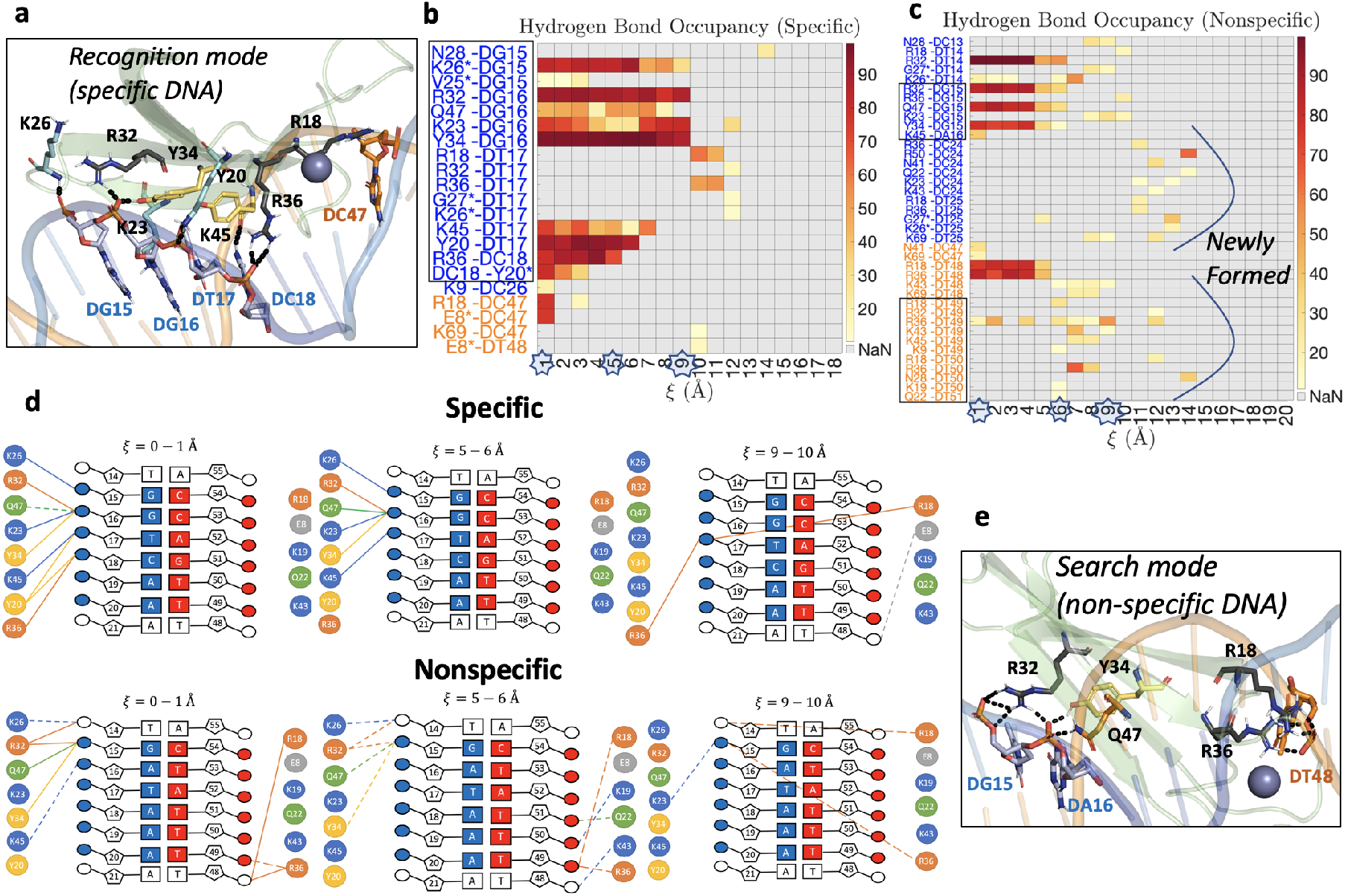
Protein-DNA interfacial hydrogen bonding (HB) interactions sampled during enforced protein dissociation from DNA (from umbrella sampling simulations). **(a)** shows the specific DNA binding system, with the protein oriented in the recognition mode. Specific HB contacts are formed, as most contacts are with the specific sequence (DG15-DC18, shown in blue), with the exception of residue R18 from a flexible loop region forming contact with the non-preferred strand (DC47, shown in orange). **(b)** and **(c)** show the average HB occupancy per umbrella window, with a preferred DNA strand (colored in blue) to associate with for the specific DNA-binding system and a non-preferred DNA strand (colored in orange). The HB contacts with the specific or nonspecific core sequence are framed. The stars on the x-axis mark the umbrella sampling windows for which the diagrams in **(d)** are drawn. **(d)** shows a schematic of the HB interactions in selected windows. The preferred DNA strand is colored in blue, the non-preferred DNA strand in orange. HBs with occupancy *>* 50% are drawn as solid lines and those with occupancy *<* 50% are drawn as dashed lines. **(e)** shows the nonspecific DNA binding system, with the protein bound to nonspecific DNA and oriented in the search mode, which lacks most of the specific contacts seen in **(a)**.

### PMF and protein dissociation kinetics from simplified spherical protein model

For the highly simplified spherical protein-DNA model system, one can calculate the free energy difference for the protein specific and non-specific DNA binding by constructing and comparing the PMFs as well from steered simulations. The Hamiltonian of the protein-DNA system coupled to a harmonic potential is *E* + 1*/*2*k*(*ξ* − *vt*)^2^, where *E* is the energy function of the protein-DNA system, and the protein center positioned at the coordinate *ξ* is constrained under the time-dependent potential centered at *vt* with force constant *k* = 4000 pN/nm, at a very low velocity of *v* = 1 Å*/μ*s. Accordingly, the simulations of steering the protein away from the DNA based on the Langevin dynamics were conducted (Figure 6a,b). One can also use the reaction coordinate *ξ* to describe the dissociation distance. As a result, the PMFs approach to flat when *ξ >* 2 nm. Thus, one can define *ξ >* 2 nm as the (micro-)dissociation. By comparing the PMFs for specific/non-specific binding, one can obtain the protein-DNA binding free energy difference ΔΔ*G*_*b*_ = 7.9 *k*_*B*_*T*. On the other hand, one can also compute the free energy difference between the protein binding to the specific site and non-specific site via the ratio of the dissociation rates, i.e., 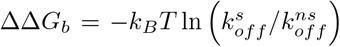, where 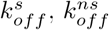 are the dissociation rates, approximately, from the specific sequence flanked by non-specific sequence (15-bp construct as in experiments) and from non-specific sequence (15-bp construct), respectively. The dissociation rate can be estimated from *k*_*off*_ = 1*/τ*_*d*_, where *τ*_*d*_ is the average dwell time of the protein from the bound DNA. Accordingly, we calculated the dwell times of the protein bound to 15-bp-sequence ‘TACGT**TTGACT**TAAT’ (with the specific core at the center), and non-specific ‘TACGTTTATGTTAAT’, repectively.

**Figure 6:**
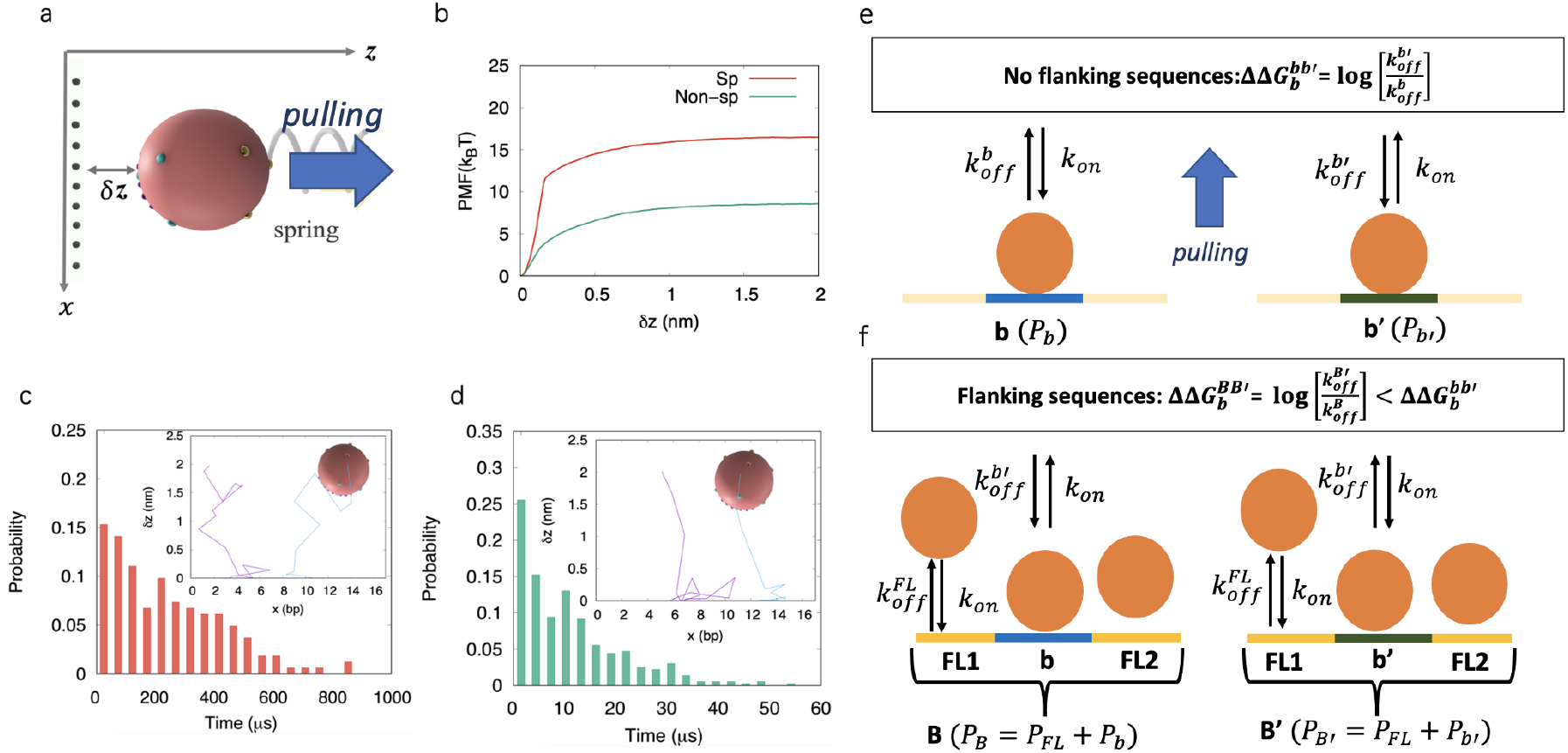
Spherical protein model comparing free energetics from steered simulation and protein-DNA dissociation kinetics, conducted for protein binding to 15-bp DNA constructs with 6-bp central specific/non-specific core DNA and flanking DNA sequences. (a) shows the schematics of steering the spherical protein by an external harmonic potential (spring). The PMFs are obtained by quasistatically pulling the protein at a center position. Note that the protein is pulled away from the 15-bp DNA with sequence ‘TACGT**TTGACT**TAAT’ with the specific sequence ‘TTGACT’ inserted at the center, and from the 15-bp non-specific DNA with sequence ‘TACGTTTATGTTAAT’, respectively. (b) shows the comparison of PMFs constructed for pulling the spherical protein away from the specific/non-specific DNAs. (c) shows the dwell-time histogram plotted for the bound population of the protein, for the protein bound to the 15-bp specific DNA. 10000 dissociation events were used. The inset shows two typical dissociation paths. (d) shows a histogram of the dwell time of the protein bound to 15-bp non-specific DNA. 10000 dissociation events were used. (e) and (f) show how the sequences flanking the binding site result in a lowered relative binding free energy for the DNA segment. (e) shows the construct with the DNA constituting of the core binding sites only, denoted as *b* (with dissociation rate 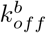) and *b*′ (with dissociation rate 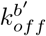). (f) shows the construct with the DNA constituting of segments *B* and *B*′ that contain the core DNA binding sites of *b* and *b*′ in the center, with flanking sites denoted by *FL* (with and approximate dissociation rate 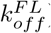). All sites are assumed to have the same association rate *k*_*on*_. The population on a site is denoted by *P*_*site*_. The corresponding difference in free energy of dissociation constructs *B* and 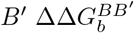 are shown to be smaller than that between site *b* and *b*′, 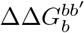.

For current purposes, we only counted the dissociation events without including re-associations. Accordingly, 10000 spontaneous dissociation events were captured from the above two 15-bp specific sequence and non-specific sequence in the simulations, respectively, similarly as being measured experimentally. The distributions of the dissociation time of the protein specifically bound to DNA (or dwell time) followed an non-exponential form (due to disparate dissociation rates from the core and from the flanking sites), with an average dwell time counted as 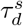. In contrast, the dwell time distribution of the non-specific core and flanking sequences were better approximated as an exponential form, with an average dwell time labeled as 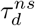 (Figure 6c-d). One should note that the respective binding free energies are likely under-estimated in the simplified spherical protein model due to a small number of protein charges used in calculating the protein-DNA electrostatic interactions, which lead to lowered dissociation times but are nevertheless expected not to affect the binding free energy difference nor the ratio of the dissociation time. The free energy difference is hence ΔΔ*G*_*b*_ = −*k*_*B*_*T* ln 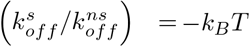 ln (1*/*25) ≈ 3.2 *k*_*B*_*T*, which turned out to be significantly smaller than that obtained from the steered simulations. The result from protein-DNA dissociation rates underestimates the free energy difference due to the center core DNA sequence (~ 6bp) being surrounded by non-specific flanking sequences, suggesting that the protein can additionally bind and unbind from the flanking DNA binding sites. On average, the flanking DNA binding sites reduce the protein-DNA binding affinity comparing to that being measured on the specific core DNA sequences.

We therefore proceeded to model the effect of the flanking DNA binding sites on the protein-DNA binding affinity by considering the scenario depicted in Figure 6e-f: we first consider DNA segments that are only constituted of the specific (*b*, with protein bound population *P*_*b*_) or nonspecific (*b*′, with population *P*_*b*_′) binding sites (see Figure 6e). At equilibrium, this gives us a difference in binding free energy 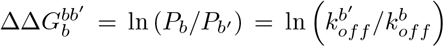. However, when *b* and *b*′ are flanked by sites *FL*1 on the left and *FL*2 on the right (see Figure 6f), with an approximate total dissociation rate 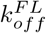 and a total protein bound population *P*_*FL*_, then the new segments *B* = *FL*1 + *b* + *FL*2 and *B*′ = *FL*1 + *b*′ + *FL*2 will have bound populations *P*_*B*_ = *P*_*b*_ + *P*_*F L*_ and *P*_*B*_′ = *P*_*b*_′ + *P*_*F L*_. The total dissociation rate will then be approximated by an average of the dissociation rates of the flanking sequences 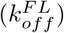 and the central sequence (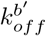 or 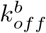), weighted by the fraction of bound protein populations to *FL* and either *b* or *b*′. We denote these fractions by 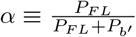 and 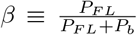. This gives us a difference in binding free energy of:

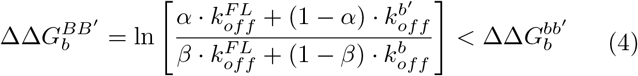

A more detailed derivation and further analysis can be found in the SI (section S-IX).

## Discussion

### The TF domain protein reorients on the DNA to switch between search (nonspecific) and recognition (specific) modes

In the facilitated diffusion model, the TF protein searches for the target DNA site by alternating between 1D diffusion along the DNA and 3D diffusion in the bulk. In order to locate the target on the DNA efficiently, it was suggested ^*b*^that the protein can adopt two conformations, which correspond to the search mode that allows for a rapid diffusion along nonspecific DNA and the recognition mode that supports stable binding of the protein to the target (specific) DNA site. Nevertheless, recent microsecond-scale MD simulations on the WRKY domain protein [19] showed that the protein bound to non-specific DNA adopts an identical conformation to that as the protein bound to the specific DNA, while the protein orientations with respect to the DNA become significantly different.

In the current study, we focused on demonstrating the binding free energetics of the WRKY domain protein on non-specific and specific DNA, respectively, to confirm that what we captured in the MD simulations well corresponded to the search and recognition modes, i.e., with significantly varied binding free energetics. Our results confirmed that the protein reorientation on the DNA indeed allows the binding free energetics to become much more stabilized from non-specific to specific DNA binding, so that the protein reorientation on the DNA can indeed support the domain protein switching from the search to the recognition mode on the DNA, without necessarily involving the protein conformational changes.

In particular, we found the relative binding free energy of the WRKY domain protein bound between non-specific and specific DNA to be 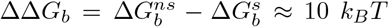 using either the umbrella sampling method or the Jarzynski equality method. It was previously predicted [18] that the transition between the TF protein search and recognition modes requires a change in the protein-DNA binding free energy of about 5-10 *k*_*B*_*T*, in agreement with current results. Current results can also be well accommodated within the two-state model, which distinguishes the search and recognition modes or states of the protein for efficient 1D search on the DNA. One should nevertheless note that current findings emphasizing protein reorientations on the DNA are mainly relevant to small globular proteins or the DNA binding domain proteins. For larger or multimeric proteins, such as LacI [38, 39, 40] or p53 [41], protein conformational changes have been indeed detected upon binding to DNA specifically vs nonspecifically.

### Protein-DNA binding free energetics in the respective search vs recognition modes

In this study, we calculated the respective protein-DNA binding free energetics of the WRKY domain protein when it is bound to the specific W-box sequence and the nonspecific DNA sequence differing slightly from the W-box (by two nucleotides). Note that such slight changes to the DNA sequences indeed lead to a significantly increased dissociation constant *K*_*d*_ (~ 80 times) with respect to the W-box sequence, as being measured experimentally [19]. In current work, to ensure the dissociation of the protein from the DNA within affordable simulation time scale for the purpose of calculating the binding free energy, we enforced the protein dissociation perpendicularly to the DNA long axis using SMD, accelerating the protein-DNA dissociation. It is important to note that the protein spontaneous dissociation is a stochastic 3D process that likely happens at millisecond time scale or over, and along various directions or paths. The enforced SMD trajectories therefore cannot represent the complete protein-DNA dissociation path space. Along the well-defined reaction coordinate *ξ* perpendicular to the DNA long axis, we indeed observe that the protein explores the horizontal degree of freedom along DNA and re-orients from time to time during dissociation, particularly in the umbrella sampling where the protein dynamics are allowed to relax for up to 200 ns for each simulation window. Consider that the horizontal protein motions average out in an ensemble of dissociation events, current samplings are able to well capture the free energy difference between the bound and unbound states of the protein with respect to the DNA, i.e, the binding free energy Δ*G*_*b*_ = *G*_*bound*_ − *G*_*unbound*_. As a sanity check, we confirmed from the literature that nonspecific protein binding had been measured to have a magnitude of 10-15 *k*_*B*_*T* for several DNA-binding proteins [42, 43], comparable to our currently measured values of around 20 *k*_*B*_*T* obtained from the Jarzynski’s and umbrella sampling methods.

Meanwhile, it is conceptually important to point out that a protein binding free energy of a magnitude of 10-20 *k*_*B*_*T* is well compatible with the diffusional search mode of the protein along DNA. In the previous work [19], where we also sampled the diffusional free energetics of the WRKY domain protein non-specifically along poly-A DNA, which show barriers ~ 2-3 *k*_*B*_*T* or a rugged free energy landscape of deviations at the thermal fluctuation level (i.e., 1-2 *k*_*B*_*T*), consistent with former experimental measurements [7]. Such a diffusional free energy profile is defined horizontally along the DNA, while the protein-DNA binding free energy Δ*G*_*b*_ is defined considering protein dissociation vertically away from the DNA. Hence, a non-specifically bound protein displaying a binding free energy of 10-20 *k*_*B*_*T* magnitude at a transient binding site can still diffuse fast along the DNA with diffusional free energy barriers 2-3 *k*_*B*_*T*. Such a diffusional search process indeed relies on shifting of hydrogen bonds at the protein-DNA interface, without involving protein dissociation that incurs at least 1-2 nm away from the DNA, vertically. Indeed, when specific target DNA sequences are located during the diffusional search process, it appears that the domain protein quickly reorients on the DNA, i.e., switched from the search to the recognition mode. Such a transition is accompanied by substantial protein-DNA binding free energy changes, e.g., from 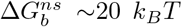 to around 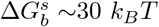, measured in current system.

### Coupled protein diffusion and protein-DNA HB dynamics during dissociation

To examine whether the protein translational and rotational degrees of freedom had been sufficiently sampled during the enforced protein dissociation, we investigated the range of protein horizontal displacements or translation degree along the DNA and the protein orientational changes. We found that in SMD simulations to construct PMF using the Jarzynski’s method, while the protein translational degrees along DNA were largely sampled, the rotational degree of protein was not well sampled yet. The limited sampling likely led to high fluctuations in the total work pulling the protein for the PMF construction. In comparison, in the umbrella sampling where the protein dynamics relaxes for a couple of hundreds of nanoseconds in each simulation window, we found that both protein horizontal displacements and rotations were appropriately sampled, and the corresponding PMF converged within about ~ 1 − 2*k*_*B*_*T*.

By investigating the HBs at the protein-DNA interface, one sees that the non-specifically-bound protein shows significantly weaker HB interactions with the DNA than the specifically bound protein, as evident from the reduced number of HB contacts in the initial bound state. In particular, the protein had about twice as many HB contacts with the 6 nucleotides constituting the specific site than those constituting the non-specific site. As a result, the protein-DNA HBs started breaking much faster upon dissociating from the nonspecific DNA (away from DNA at 5-6Å) than from the specific DNA (away at 9-10 Å). Interestingly, the protein dissociated from the nonspecific DNA started re-forming contacts with both DNA strands during the dissociation. In addition, the domain protein was observed to randomly probing the DNA surface, consistent with what would be expected for the TF protein in the search mode.

On the other hand, the specific complex maintained the initial HB contacts at the protein-DNA interface, mainly on the preferred DNA strand, as the protein core domain (including the WRKY motif) interacts exclusively with the specific W-box sequence. After the initial breaking of the HBs, very few new protein-DNA HB contacts re-form (~ 4 times fewer than the number of contacts re-formed in the nonspecific system). Among those new protein-DNA HB contacts formed, only two were with the non-preferred strand. The tighter binding of the protein to the specific DNA and the strong DNA sequence preference are also consistent with what would be expected from a protein in the recognition mode.

### Toy model and analyses reveal potential impacts of flanking DNA sequence on measurements of protein-DNA affinity

The protein-DNA binding free energy difference calculated in current simulation studies between specific and nonspecific DNA complexes was about 10*k*_*B*_*T*, obtained using either the Jarzynski’s equality or the umbrella sampling methods. In contrast, the experimentally obtained free energy difference was estimated ~ 4.2 *k*_*B*_*T*, according to the measured difference of the protein dissociation constants for the non-specific and specific DNA complexes, with DNA constructs of 15-bp [20].

Considering that the WRKY domain protein binding site occupies only about 6 bps on the DNA, we investigated a potential source of this discrepancy in ΔΔ*G*_*b*_ in a highly simplified spherical protein model with a 15-bp DNA in complex, which is the same length as in the experimental measurements [20]. To compare, 6-bp specific and non-specific core DNA sequences were designated at the center of the 15-bp DNA. In one simulation setup, the spherical protein was steered vertically away from the central DNA binding site through a spring force, similar to SMD simulation setup. In the other setup, the initially bound protein was allowed to diffuse or dissociate spontaneously from DNA using the stochastic simulations, and the subsequent dissociation events were recorded. The latter method is to mimic the experimentally measured real protein-DNA dissociation dynamics, capturing potential effects that DNA sequences flanking the binding site can support both the protein binding and the dissociation. It is worth noting that both *in vitro* and *in silico* analyses have shown that DNA sequences flanking the DNA core motifs indeed influence the affinity of the TF to the motif [44, 45, 46, 47] by altering the 3D structure and the flexibility of the DNA. However, to our knowledge, no kinetic models have been developed so far to account for such effects on protein-DNA binding affinities.

The results from the first setup, i.e., the steered simulations show that 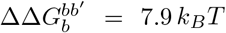 between the DNA constructs with the non-specific and specific DNA core (b’ and b), while that from the stochastic simulations of the spontaneous protein dissociation kinetics on the 15-bp DNA actually suggest a reduced free energy difference as 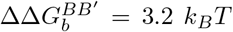. The result appears consistent with the discrepancy observed between our all-atom MD simulations and experimentally estimated values, with 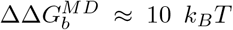 (comparable to 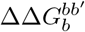), and 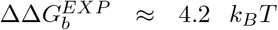 (comparable to 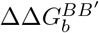). The results also suggest that the DNA sequences flanking the central binding site do have significant impact on measuring the protein-DNA dissociation kinetics or binding affinity. Indeed, the protein can bind to both the central binding site and the sites flanking it (e.g. via diffusion from the central site). If one considers that the protein has a lower binding affinity to the non-specific flanking DNA sequences than to the central specific DNA sequence, then the protein can bind and then unbind from these sites with various unbinding rates *k*_*off*_. Consequently, the dissociation constant is influenced by all possible dissociation events or kinetics from central and the flanking DNA sites. Indeed, an overall dissociation rate, though not well defined as for a Poisson process, can be approximated by averaging the protein dissociation rates from the central binding sites and flanking sequences, weighted by the populations at corresponding sites. Since the dissociation rate of the flanking sequences are larger than that of the central binding site with specific DNA sequences, the overall approximate or averaged dissociation rate will then be larger than that of the central binding site alone. As a result, effectively, the relative binding free energy between the non-specific and specific DNA construct is measured lower than that for the central site in the absence of protein binding to the flanking DNA sequences. That is, 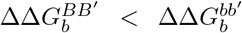, which can possibly explain 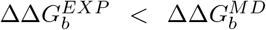. It is worth noting that both *in vitro* and *in silico* analyses have shown that DNA sequences flanking the DNA core motifs indeed influence the affinity of the TF to the motif [44, 45, 46, 47] by altering the 3D structure and the flexibility of the DNA. However, to our knowledge, no kinetic models have been developed so far to account for such effects on protein-DNA binding affinities. Further experimental studies with designs of the protein binding to various lengths of DNA constructs are needed, both to improve the kinetic model and to better elucidate the impact of flanking sequences on the protein-DNA binding affinities.

## Data Availability

Computing codes available upon request

## Acknowledgements

J.Y. has been supported by the Center for Multiscale Cell Fate Research of UCI (University of California, Irvine) via NSF (CMCF) Division of Mathematical Sciences Grant 1763272, Simons Foundation Grant 594598, and start-up funding from UCI. C.M. has been supported by an NSF grant, DMS1763272, and CMCF fellowship from the Simons Foundation (594598).

