## Supplemental Materials for "Non-specific vs specific DNA binding free energetics of a transcription factor domain protein for target search and recognition"

### Supplementary Materials

#### S-I Equilibrium properties of the specific and nonspecific binding systems

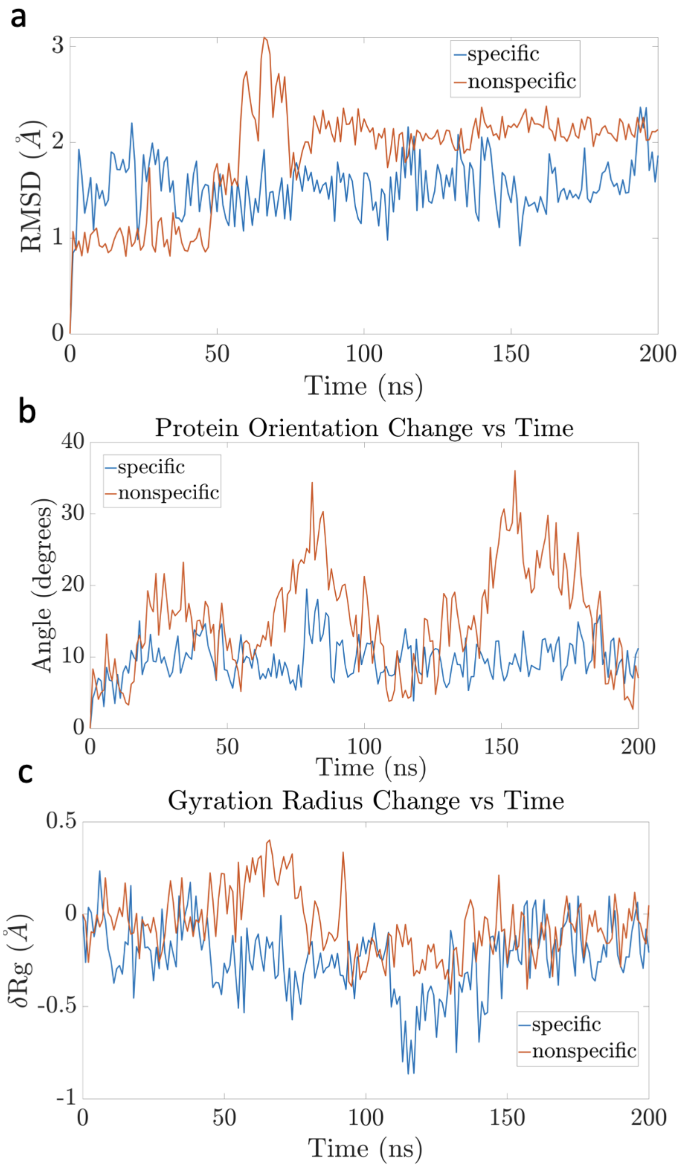


Figure S1: Equilibrium properties of the protein bound to both specific (blue) and nonspecific (red) DNA. The flexible peripheral loops were excluded from the measurements (protein residues 8-15 and 63-69). **(a)** shows the RMSD of the protein in both specific and nonspecific systems, stabilized starting about 80 ns. **(b)** shows the average orientational changes with respect to the initial frame of the four beta strands of the protein in both systems, with the average orientational change equal to $16.6° \pm 7.9°$ for the nonspecific system and $10.1° \pm 2.8°$ for the specific system. **(c)** shows the change in gyration radius $\delta Rg$ of the protein in both systems, with an average change of -0.06±0.16 Å for the specific system and -0.25±0.19 Å for the nonspecific system.

### S-II RAMD to determine protein dissociation paths

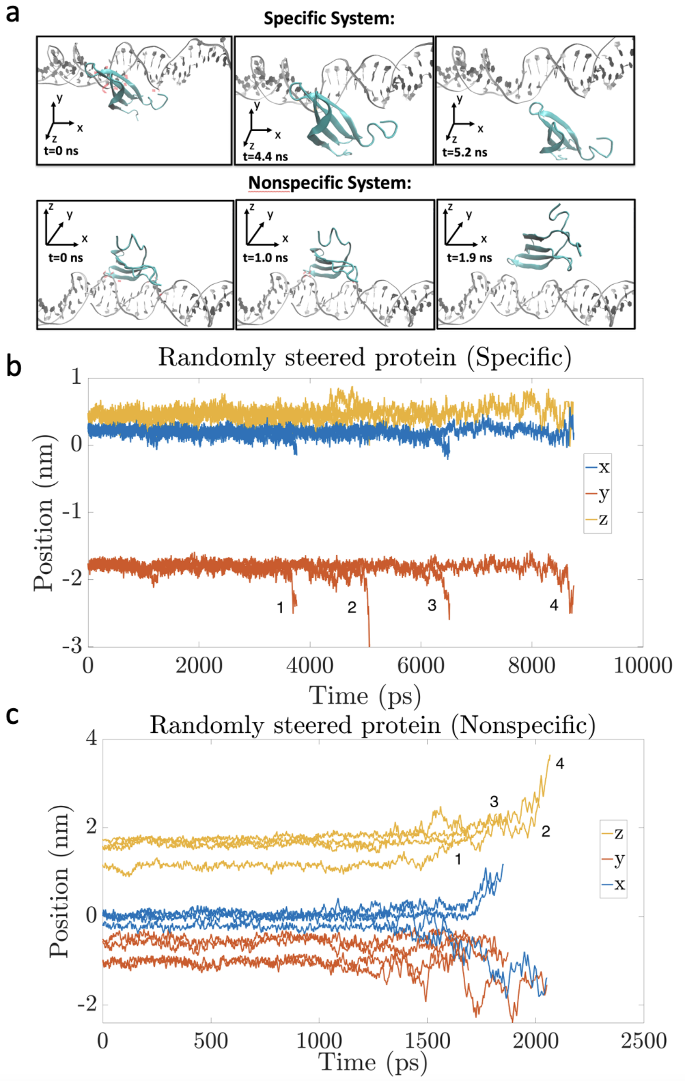


Figure S2: Preferred direction of protein dissociation for both specific and nonspecific DNA binding systems. The reaction coordinates were chosen by finding the optimal dissociation path using randomly accelerated MD (RAMD). Five trajectories were run for each system. The center of mass of the protein is steered in a random direction with a constant force k=2000 kJ/mol. If the protein COM does not move by a distance larger than 0.025 nm in 100 fs, the direction is updated to another random one. **(a)** shows snapshots of a sample RAMD trajectory for each system. **(b)**,**(c)** show the protein COM displacement in all 3 directions for 4 RAMD trajectories for both specific and nonspecific systems, respectively. We find that the preferred direction of dissociation is the -y direction for all 4 trajectories in the specific DNA system, and +z direction for 3 out of 4 trajectories in the nonspecific system (the other being in the -y z plane)

#### S-III Jarzynski's Method: Protocol

The steering protocol is illustrated in Fig.S3. "Forward pulling" (blue arrows) refers to the steering of the protein away from the DNA (-y direction specific, z-direction nonspecific) and "backwards pulling" refers to steering the protein in the opposite direction, towards the DNA. The protein was pulled, equilibrated, then pulled again. Two different sets of forward and backward pulling were conducted and can be summarized as follows:

- Steer the protein from $\xi = 1 \to 0$, $1 \to 2$, $3 \to 2$, $3 \to 4$, $8 \to 4$, $8 \to 12$, $16 \to 12$, $16 \to 20$ (in units of Å).
- At each starting point, equilibrate the system for 2 ns.
- Allow the forward and backward pulling to overlap over a region of width 0.5 A, and take the average work applied in that region
- Repeat 5 times for each segment
- Repeat the above procedure, this time with $\xi = 0 \to 1$, $2 \to 1$, $2 \to 3$, $4 \to 3$, $4 \to 8$, $12 \to 8$, $12 \to 16$, $20 \to 16$ (in units of Å).

With this procedure we end up with 10 trajectories per segment from which we can get the corresponding PMF.

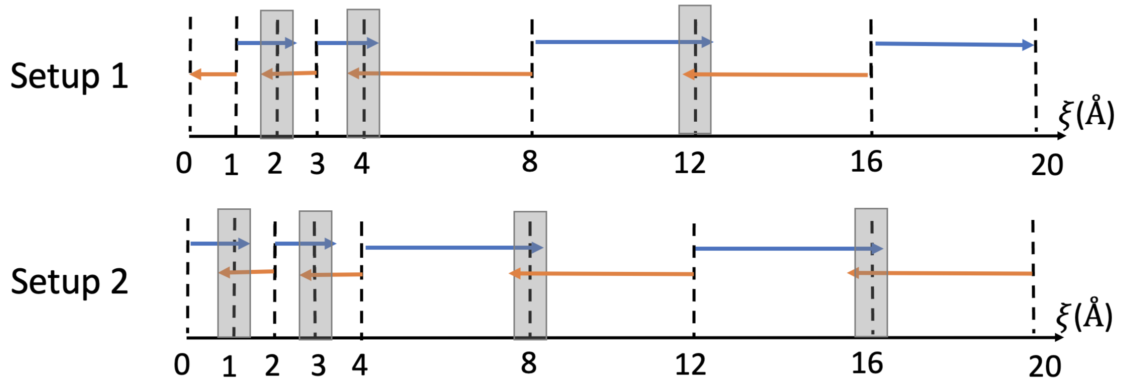


Figure S3: Protocol for steering the COM of the protein to construct the PMF using Jarzynski's equality. Two different sets of forward and backward pulling were implemented, corresponding to setup 1 and setup 2 trajectories in Fig.3a. The blue arrows represent forward pulling and orange arrows the backwards pulling. The overlap region is shown in grey. The dashed lines indicate the start and end points the pulling. At every starting point the structure is equilibrated for 2 ns.

### S-IV Umbrella sampling method

#### S-IV.1 HB occupancy along the dissociation path obtained from SMD

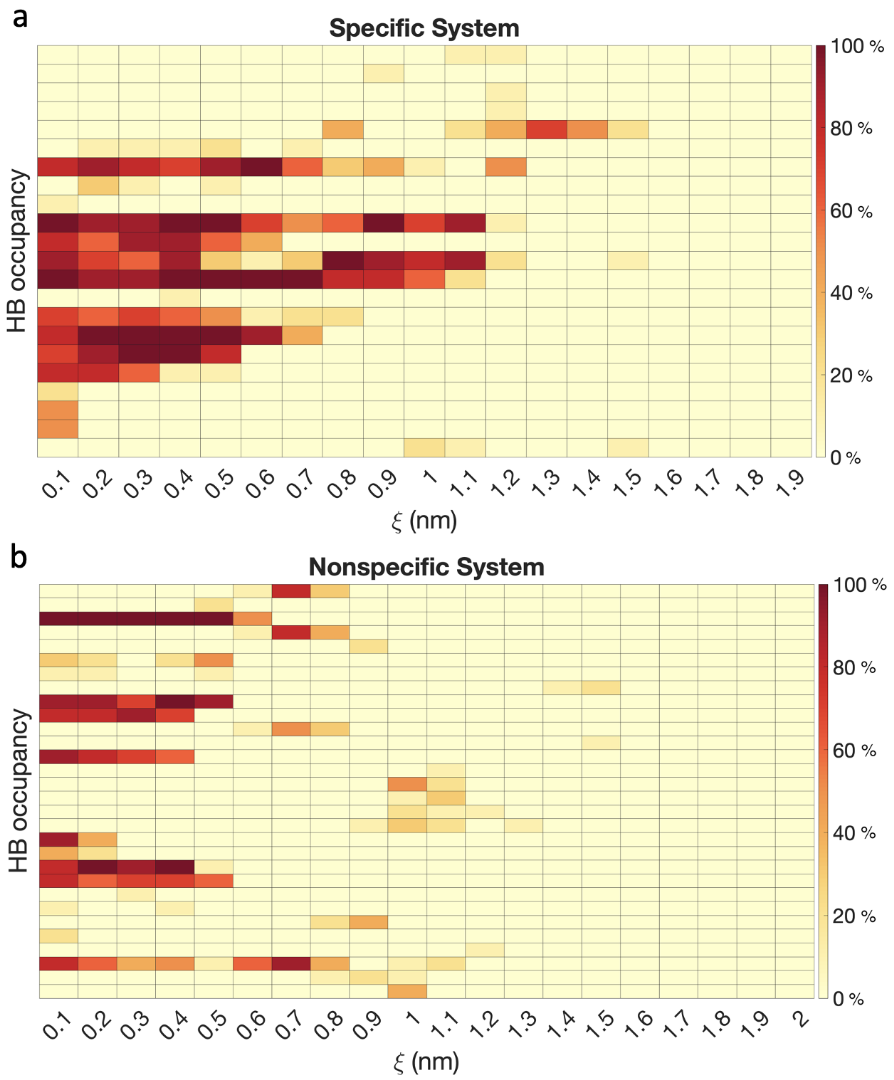


Figure S4: HB occupancy along the dissociation path obtained from SMD. The heatmaps show the average occupancy (in percentages) of HB contacts per residue pair over the entire pulling trajectory for both the specific **(a)** and nonspecific **(b)** systems. The HB distance cutoff was chosen to be 0.35 nm, and the angle cutoff 30°. HB contacts gradually decreased and were completely broken starting $\xi = 1.6$ nm, which indicates that the protein has dissociated microscopically. This serves as a sanity check to make sure the trajectories properly involve full dissociation of the protein.

#### S-IV.2 Umbrella Sampling PMF convergence over time

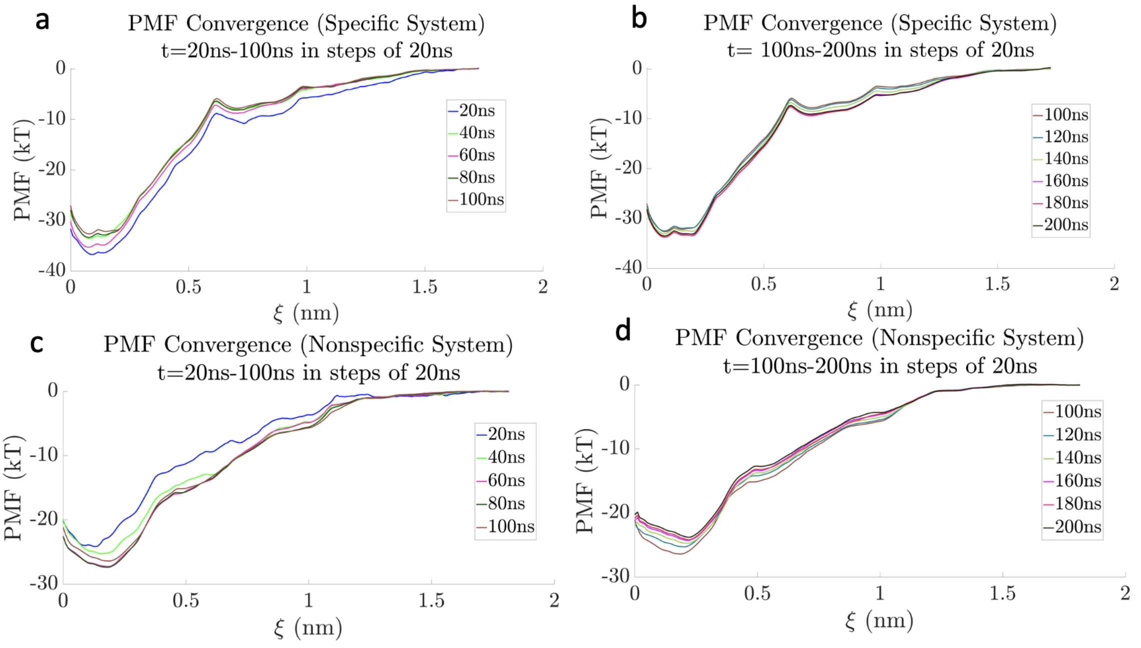


Figure S5: PMFs from umbrella sampling method for protein dissociation from DNA change when averaging over a longer cumulative trajectory time for both specific and nonspecific systems. **(a),(b)** show the PMFs from 0-100ns, and **(c),(d)** those from 100-200ns were plotted separately. By around 150ns, the PMF for both the specific and nonspecific systems converged within 1.5 $k_BT$.

#### S-V Langevin dynamics of the spherical protein

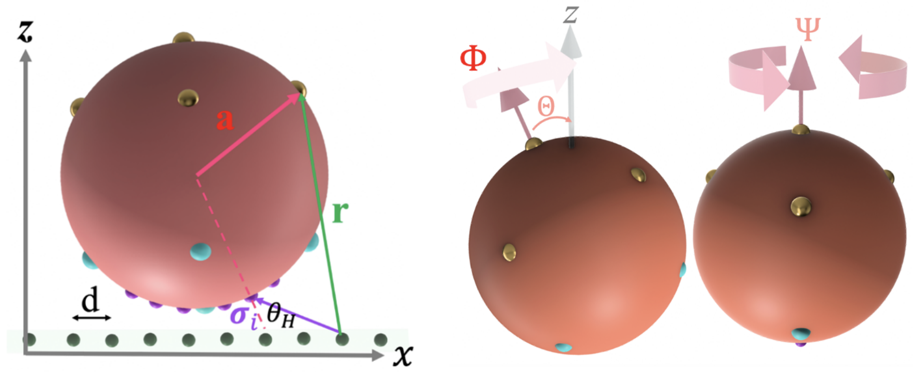


Figure S6: The simplified spherical protein-DNA binding system. Left: The spherical protein model toy model, with the DNA-protein interaction sites shown. The effective positive charges and negative charges are indicated by cyan and coral dots, respectively. The hydrogen bond (HB) interaction sites (violet dots) are mounted on the positively charged hemisphere. Binding sites on DNA are embedded in a tube (along the x-axis) with a diameter of 0.2 nm and aligned on the axis of the tube separated individually by d=0.34 nm in distance. Vector **r** stands for the vector between the i-th DNA binding site and an interaction site on the protein, and the radial vector **a** in spherical coordinates, (i.e., $|\mathbf{a}|, \theta, \phi$). Bond angle $\theta_H$ between a HB interaction site on the protein and a site of DNA is measured between the radial direction (dashed line) of the HB interaction site on the sphere and the direction (soiled line) of the pair-interaction, i.e., $\cos\theta_H = -a \cdot \sigma_i$. Right: the rotational degrees of freedom of the spherical protein. An axial direction (red arrow) represents the inverse direction of the electrostatic polarization of the protein (here from positive to negative), which can be described by spatial angles Θ (relative to the z-axis) and Φ (relative to the x-axis) with an additional Ψ for protein spinning about its axis.

### S-VI Simplified spheric protein-DNA model construction

The domain protein can be approximated by a charged sphere (Fig.S6). Discretized positive and negative charges were placed on the surface of the spherical protein under overall neutrality. The protein-binding sites on DNA are represented by linearly aligned points with spacing of 0.34 nm. In addition, the hydrogen bond (HB) interaction are represented by the pair-interaction between the sites on the protein hemisphere and the sites on DNA. The protein-DNA electrostatic interactions screened by ions can be approximated by the Debye-Huckel potential, and the HB interaction can be described by the Morse potential depending on the bond angle. Thus, the energetics between the spherical protein and the linear DNA is:

$$E(x,y,z,\Theta,\phi,\Psi) = U_{ele}+U_{HB} = \sum_{i,j} \frac{q_i Q_{p,I}}{4\pi\epsilon_0\epsilon_r r}\exp[-\kappa r]+\sum_{i,j}\mathcal{E}_{HB,J}\left[\left(e^{-\alpha(\sigma_i-r_0)}-1\right)^2-1\right]\cos\theta_H \tag{S-1}$$

where the coordinates $x, y, z$ describe the center of mass of the protein, and $\Theta$,$\Phi$ and $\Psi$ describe the orientation of the spherical protein, $q_i = -2e$ is the effective charge of one pair of nucleotides on the $i$-th site on DNA, $Q_{p,I}$ is the effective charge of the $I$-th charged site on the spherical protein, $\epsilon_{r}is$ the dielectric constant, $\mathbf{r}$ the distance between the $i$-th DNA binding site and the charged site on the protein (Fig.S6, right) and $\kappa$ the inverse Debye length, characterizing the strength of the electrostatic screening, $E_{HB,J}$ is the depth of the Morse potential of the $J$-th HB site on protein, $\alpha$ is the inverse decay length, $r_0$ specifies the radius of the repulsive core, $\sigma_i$ the distance between the $i$-th DNA binding site and the HB site on the protein , and $\theta_H$ is the bond angle.

As for the Langevin dynamics of this spherical protein model, we set the radius of the spherical protein to 1.5nm. 10 discretized pointed-like charges of $Q_p = \pm 0.55e$ each are placed on the surface of the protein. The effective charge of one pair of nucleotides on the i-th site of DNA is $q_i = -2e$. The inverse Debye length $\kappa$ characterizing the strength of the electrostatic screening. The strength $\mathcal{E}_{HB,J}$ is sequence-dependent, where $J$ takes 1 to 6, denoting 6 HB interaction sites on the protein. We can define the strength of HB interaction as follows. $\mathcal{E}_{HB,1} = 7.2\ k_BT$ if the 1st site binds to T; $\mathcal{E}_{HB,2} = 7.2\ k_BT$ if the 2nd site binds to T; $\mathcal{E}_{HB,3} = 7.2\ k_BT$ if the 3rd site binds to G; $\mathcal{E}_{HB,4} = 7.2\ k_BT$ if the 4th site binds to A; $\mathcal{E}_{HB,1} = 7.2\ k_BT$ if the 5th binds to C; $\mathcal{E}_{HB,2} = 7.2$ $k_BT$T if the 6th site binds to T; and $\mathcal{E}_{HB,J} = 0.8\ k_BT$, otherwise. This rule charactizes the specific sequence'TTGACT'. The inverse decay length is $\alpha$, and the repulsive core radius $r_0$.
To exclude the effects of the mobility of DNA, we freeze the DNA, Thus, the COM (center of mass) of the protein evolves under the full potential $E(x, y, z, \Theta, \phi, \Psi)$,

$$\dot{\boldsymbol{R}}_{COM} = -\frac{1}{\zeta}\nabla_{RCOM}E + \sqrt{\frac{2k_BT}{\zeta}}\boldsymbol{w} \tag{S-2}$$

where $\zeta = 6\pi\eta a$ is the friction to the protein in solution, $k_B$ is the Boltzmann constant and $\boldsymbol{w}$ is the Gaussian noise with zero mean and unit variance. The time-step is 1 pico-second.
The 3-dimensionanl rotations caused by the thermal fluctuations and interactions can be divided into a spatial (2-dimensional) displacement of the unit vector A (axial vector of the protein), $\Delta\Omega_A$, and the 1-dimensional spin about $A$, $\Psi$ (Fig.S6).

$$\begin{aligned}\boldsymbol{A}\times\dot{\boldsymbol{\Omega}}_A|\boldsymbol{a}| &= -\frac{1}{\zeta_{rot}}\sum_i(\boldsymbol{a}\times F_i)\times\frac{\boldsymbol{A}}{|\boldsymbol{a}|}+\sqrt{\frac{2k_BT}{\zeta_{rot}}}(\boldsymbol{A}\times\boldsymbol{e}_\Theta w_1+\boldsymbol{A}\times\boldsymbol{e}_\Phi w_2)\\ \boldsymbol{A}\cdot\dot{\boldsymbol{\Psi}}|\boldsymbol{a}| &= -\frac{1}{\zeta_{rot}}\sum_i(\boldsymbol{a}\times F_i)\times\frac{\boldsymbol{A}}{|\boldsymbol{a}|}+\sqrt{\frac{2k_BT}{\zeta_{rot}}}w_3\end{aligned} \tag{S-3}$$

where $\zeta_{rot} = 8\pi\eta a$ is the friction due to rotation of the protein in solution, $w_1$,$w_2$ and $w_3$ are Gaussian noises with zero mean and unit variance.

#### S-VII Jarzynski's Method: Distribution of the work applied during protein dissociation

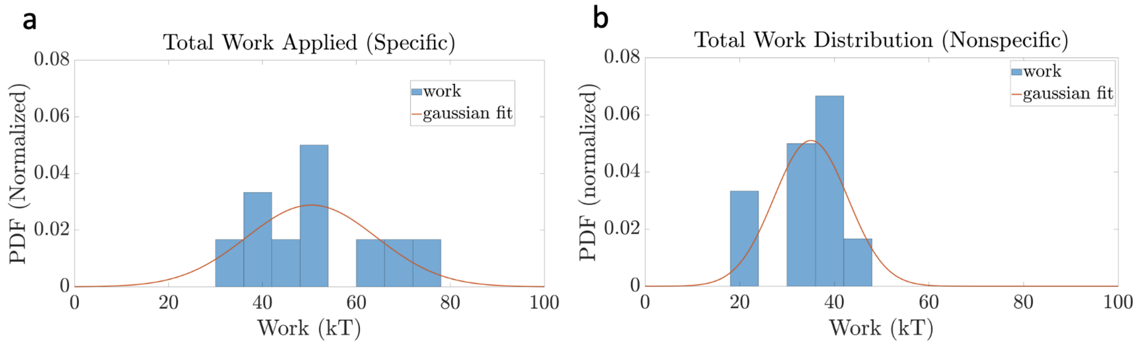


Figure S7: Distribution of the total work applied in protein dissociation from the DNA for the PMF construction using Jarzynski's equality. **(a)** and **(b)** show the total work distribution (average taken for the trajectory corresponding to the reaction coordinate $\xi \geq 1.5$ nm), which deviate from a gaussian, particularly for the specific DNA binding system.

#### S-VII.1 Jarzynski's method: Convergence of PMFs

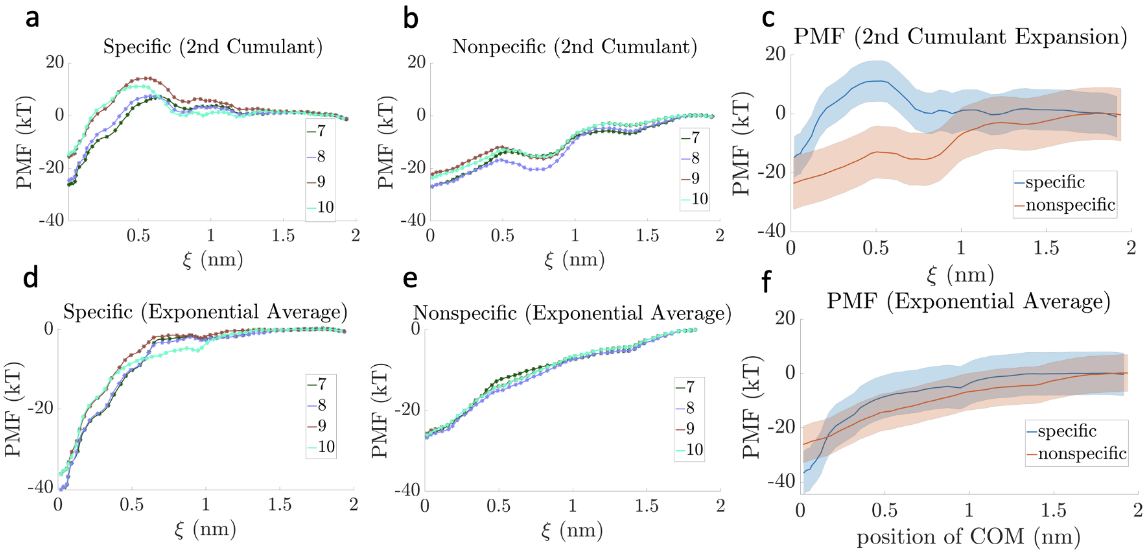


Figure S8: PMF constructed from the Jarzynski equality method convergence by adding one trajectory at a time (last 4 trajectories, trajectories 7-10 shown). **(a)** and **(b)** show the convergence of the PMFs obtained from the second cumulant expansion (main text, Eq.(2)) for both systems, with the corresponding PMFs shown in **(c)**. The PMFs show that the specific DNA binding system has a lower energy barrier than the nonspecific one. **(d)** and **(e)** show the convergence of the PMFs obtained from taking the exponential average (main text, Eq.(1)) when each of the trajectories is added at a time. Specific DNA binding system converges within 4 $k_BT$, nonspecific one within 2 $k_BT$. The corresponding PMFs are shown in **(f)**, with the PMF standard deviations shown as the shaded regions.

#### S-VIII Umbrella Sampling: Distribution of the reaction coordinate in each umbrella window

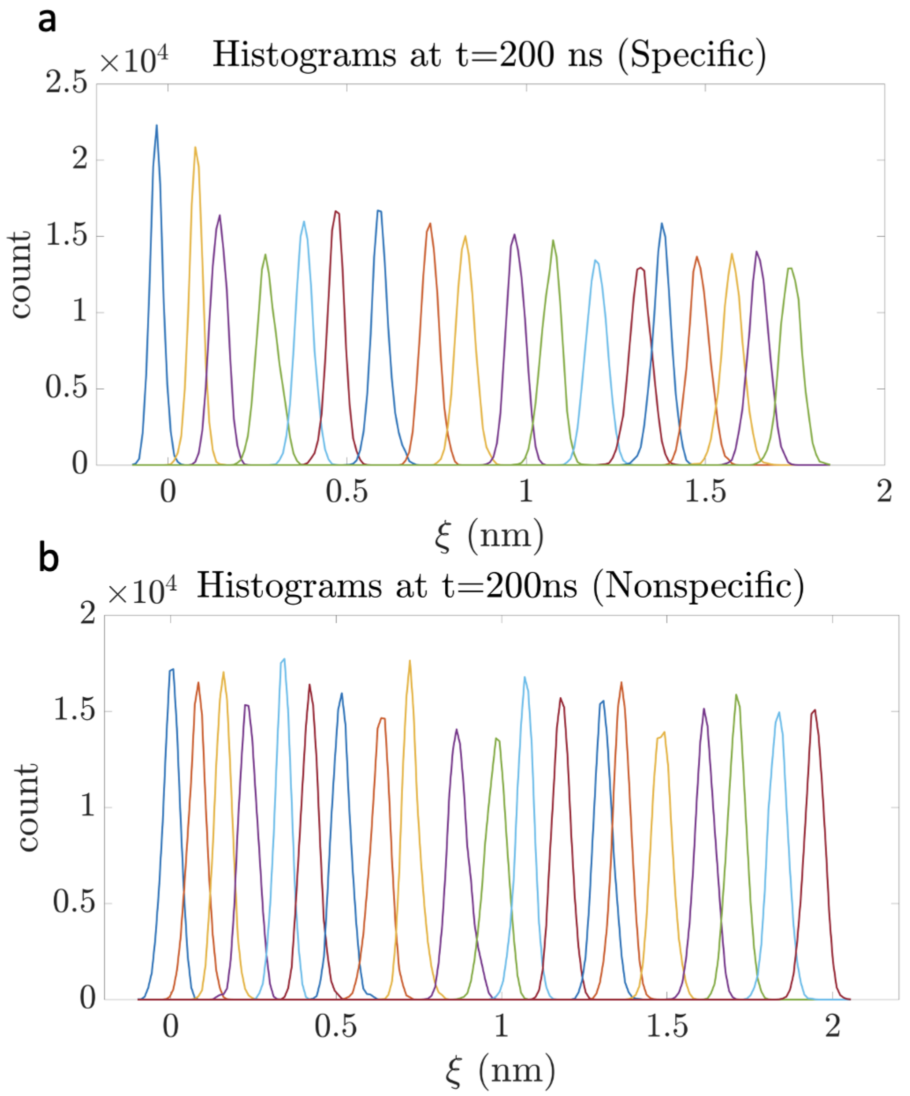


Figure S9: Histograms obtained from umbrella sampling simulations showing the distribution of the reaction coordinate sampled over each window for both specific **(a)** and nonspecific **(b)** DNA binding systems of the WRKY. The reaction coordinate $\xi$ is measured as the center of mass of protein displaced with respect to the initial position, or say, along the vertical dissociation path from DNA. A force constant of $k = 3000$ kJ.mol$^{-1}$nm$^{-2}$ was used to keep the COM of the protein fixed along $\xi$.

### S-IX Potential relationship between experimental and computational binding affinities

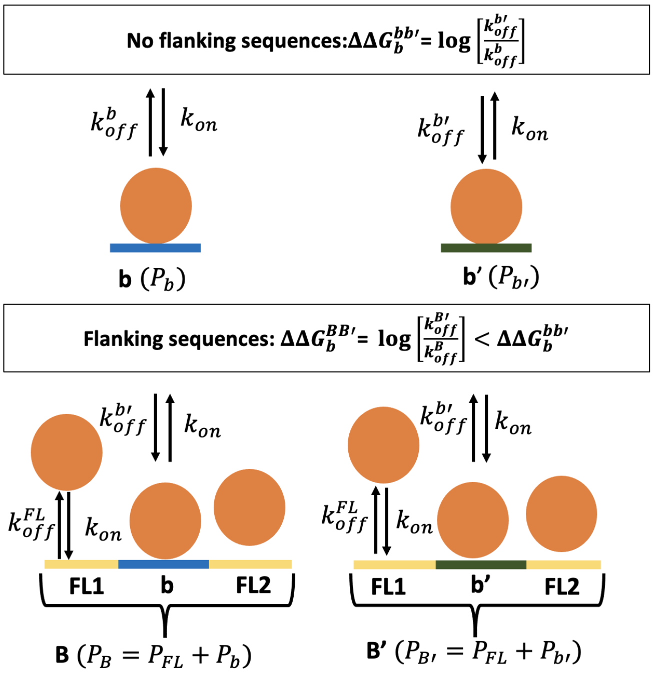


Figure S10: Sequences flanking the binding site result in a lowered relative binding free energy for the DNA segment. Top: construct with the DNA constituting of the core binding sites only, denoted as $b$ (with dissociation rate $k_{off}^{b}$) and $b'$ (with dissociation rate $k_{off}^{b'}$). Bottom: construct with the DNA constituting of segments $B$ and $B'$ that contain the core DNA binding sites of $b$ and $b'$ in the center, with flanking sites denoted by $FL$ (with and approximate dissociation rate $k_{off}^{FL}$). All sites are assumed to have the same association rate $k_{on}$. The population on a site is denoted by $P_{site}$. The corresponding difference in free energy of dissociation constructs $B$ and $B'$ $\Delta\Delta G_b^{BB'}$ are shown to be smaller than that between site $b$ and $b'$, $\Delta\Delta G_b^{bb'}$.

### S-X Flanking DNA sequence impacts to protein dissociation kinetics

In this section, we proceed to derive a kinetic model that accounts for the impact of flanking DNA sequences in the protein dissociation kinetics, which will consequently impact the protein-DNA binding free energy measurements.

The dissociation constant is given by:

$$K_d = \frac{[prot][DNA]}{[prot.DNA]} = \frac{k_{off}}{k_{on}} \tag{S-4}$$

Where $[prot]$, $[DNA]$ and $[prot.DNA]$ represent respectively the concentration of unbound free protein, the concentration of unbound free DNA and the concentration of protein-DNA complexes. $k_{on}$ is the association rate constant ($M^{-1}s^{-1}$) , i.e, the number of protein-DNA complexes forming in 1 Molar of solution per second, and $k_{off}$ is the dissociation rate constant ($s^{-1}$), i.e, the number of complexes that dissociate per second. The corresponding change in Gibbs free energy is then given by:

$$\Delta G_b = k_B T \ln\left(\frac{K_d}{c_0}\right) \tag{S-5}$$

Where $k_B$ is the Boltzmann constant, $T$ the temperature of the system and $c_0$=1 mol/L is the standard reference concentration.
Now consider two separate DNA segments in a solution containing proteins, $b$ and $b'$, corresponding respectively to the specific and nonspecific binding sites. The corresponding dissociation rates are then $k^b_{off}$ and $k^{b'}_{off}$. We can assume that the protein binding events are independent, making $k_{on}$ the same for each segment. We can also denote the population of proteins bound to $b$ by $P_b$, and those bound to $b'$ by $P_{b'}$. The setup is shown in diagram Fig.S10. At equilibrium, the flux of proteins dissociating from respective segments are equal, giving us the relationship:

$$\begin{aligned} P_b \times k^b_{off} &= P_{b'} \times k^{b'}_{off} \\ \Delta\Delta G^{bb'}_b &= \ln \frac{k^{b'}_{off}}{k^b_{off}} \end{aligned} \tag{S-6}$$

Where $\Delta\Delta G^{bb'}_b$ is the free energy difference between the specific and nonspecific binding sites.
Now consider the segments $B$ and $B'$ constituted of three sub-segments $(FL1, b, FL2)$ and $(FL1, b', FL2)$, where $b$ and $b'$ are the central specific and nonspecific binding sites, respectively, and $FL1, FL2$ the left and right flanking sequences, to which the protein has a much lower affinity than to a specific DNA binding site (or equivalent to non-specific binding site). The flanking sites have a dissociation rate $k^{FL}_{off}$ (as an approximation) and the specific and nonspecific binding sites $k^b_{off}$ and $k^{b'}_{off}$ respectively. The populations are then given by $P_B = P_{FL} + P_b$ and $P_{B'} = P_{FL} + P_{b'}$. This gives us:

$$\begin{aligned} P_B \cdot k^B_{off} &= (P_{FL} + P_b) \cdot k^B_{off} = P_{FL} \cdot k^{FL}_{off} + P_b \cdot k^b_{off} \\ P_{B'} \cdot k^{B'}_{off} &= (P_{FL} + P_{b'}) \cdot k^{B'}_{off} = P_{FL} \cdot k^{FL}_{off} + P_{b'} \cdot k^{b'}_{off} \end{aligned} \tag{S-7}$$

Note that $k^B_{off}$ and $k^{B'}_{off}$ are defined approximately (not single kinetic event). At equilibrium, the fluxes are equal, and the free energy difference between the specific and nonspecific binding sites are now given by:

$$\Delta\Delta G^{BB'}_b = \ln \frac{k^{B'}_{off}}{k^B_{off}} = \ln \frac{\alpha \cdot k^{FL}_{off} + (1-\alpha) \cdot k^{b'}_{off}}{\beta \cdot k^{FL}_{off} + (1-\beta) \cdot k^b_{off}} \tag{S-8}$$

where $\alpha == \frac{P_{FL}}{P_{FL}+P_{b'}}$ and $\beta == \frac{P_{FL}}{P_{FL}+P_b}$ represent the fraction of proteins bound to the flanking sequences in $B'$ and $B$, respectively. Since $k^{FL}_{off}$ is comparable to that of non-specific DNA binding sites, and $k^b_{off}$ is for the specific DNA binding site, $k^{FL}_{off} \gg k^b_{off}$, Eq.S-6 and Eq.S-8 then imply that $\Delta\Delta G^{BB'}_b < \Delta\Delta G^{bb'}_b$.

This equation can be further simplified by assuming that the flanking sites are nonspecific binding sites with the protein binding affinity comparable to $b'$. Then $k_{off}^{FL} \sim k_{off}^{b'} \gg k_{off}^{b}$. Then Eq. S-8 becomes:

$$\Delta\Delta G_b^{BB'} = \ln \frac{k_{off}^{b'}}{\beta \cdot k_{off}^{b'} + (1-\beta) \cdot k_{off}^{b}} = \ln \frac{\frac{k_{off}^{b'}}{k_{off}^{b}}}{\beta \cdot \frac{k_{off}^{b'}}{k_{off}^{b}} + (1-\beta)} \tag{S-9}$$

Experimentally, $\Delta\Delta G_b^{BB'} \approx 5\ k_B T$. Now suppose that the difference in binding free energy between the specific and nonspecific binding sites is around 10 $k_B T$ (from our MD simulation results). Then one sets $\Delta\Delta G_b^{bb'} = \ln k_{off}^{b'}/k_{off}^{b} = 10\ k_B T$, so that $k_{off}^{b'}/k_{off}^{b} \approx 10^4$. Additionally, since $b$ is the specific binding site, we can assume that the proteins will most likely bind to $b$, giving us $P_b \gg P_{FL}$, which implies $(1-\beta) \approx 1$ and $\beta \ll 1$. Putting it all together, we finally get:

$$\Delta\Delta G_b^{BB'} = \ln \frac{10^4}{\beta 10^4 + 1} = 5\ k_B T \tag{S-10}$$

This gives us $\frac{10^4}{\beta 10^4+1} \approx 100 \rightarrow P_{FL}/P_b = \beta \approx 0.01$, implying that only 1% of the proteins need to be bound to the flanking sequences for $\Delta\Delta G_b^{BB'}$ to be significantly lower than $\Delta\Delta G_b^{bb'}$.

In other words, when the specific binding site is flanked by nonspecific, lower affinity sites, effective protein dissociation rate or kinetics enhances. This can explain why the experimentally obtained binding free energy $\Delta\Delta G_b^{exp}$ influnced by flanking DNA binding sites on the 15-bp DNA is lower than that obtained from MD simulations designed for calculating for the central DNA binding site (6-bp) as $\Delta\Delta G_b^{MD}$. While this model is simplified and hence does not recover the exact relationship between $\Delta\Delta G_b^{exp}$ and $\Delta\Delta G_b^{MD}$, it does provide valuable insight that can explain the difference between these relative binding free energies.
